# *PKD1* and *PKD2* mRNA cis-inhibition drives polycystic kidney disease progression

**DOI:** 10.1101/2022.02.23.479396

**Authors:** Ronak Lakhia, Harini Ramalingam, Chun-Mien Chang, Patricia Cobo-Stark, Laurence Biggers, Andrea Flaten, Jesus Alvarez, Tania Valencia, Darren P. Wallace, Edmund Lee, Vishal Patel

## Abstract

Autosomal dominant polycystic kidney disease (ADPKD), among the most common human genetic conditions and a frequent etiology of kidney failure, is primarily caused by heterozygous *PKD1* mutations. Kidney cyst formation ensues when the *PKD1* dosage falls below a critical threshold. However, no framework exists to harness the remaining allele or reverse *PKD1* decline. Here, we show that mRNAs produced by the noninactivated *PKD1* allele are cis-repressed *via* their 3’-UTR miR-17 binding element. Eliminating this motif (*Pkd1*^Δ17^) improves mRNA stability, raises Polycystin-1 levels, and alleviates cyst growth in cellular, *ex vivo*, and mouse PKD models. Remarkably, *Pkd2* is also autoinhibited *via* its 3’-UTR miR-17 motif, and *Pkd2*^Δ17^-induced Polycystin-2 derepression partly compensates and retards cyst growth in *Pkd1*-mutant models. Moreover, acutely blocking *Pkd1/2* cis-inhibition, including after cyst onset, attenuates murine PKD. Finally, *PKD1*^Δ17^ or *PKD2*^Δ17^ alleles revert cyst-pathogenic sequala in patient-derived primary ADPKD cultures. Thus, evading 3’-UTR cis-interference and enhancing *PKD1/2* mRNA translation is a potentially mutation-agnostic ADPKD-arresting approach.

## INTRODUCTION

An estimated 12.5 million people worldwide suffer from autosomal dominant polycystic kidney disease (ADPKD), making it among the most common monogenetic conditions known to humankind. A clinical hallmark of ADPKD is the relentless growth of innumerable fluid-filled cysts in the kidneys, which replace the normal parenchyma and over decades cause massive bilateral kidney enlargement and renal failure^1^. ADPKD occurs because of heterozygous, loss-of-function mutations in *PKD1* (~78% of cases) or *PKD2* (~15% of cases). The classical hypothesis for cyst initiation is that in addition to a germline inactivating mutation in one allele of the PKD gene, there is somatic inactivation (referred to as ‘second hit’) in the other allele, causing a complete loss of polycystin expression in the cell. However, in recent years, several lines of evidence support the gene dose threshold as a mechanism involved in cystogenesis^2,3^. This hypothesis posits that complete *PKD1* loss is not necessary, but rather cystogenesis ensues if the *PKD1* dosage falls below a critical threshold. Supporting the gene dosage model,‘second hit’ mutations do not appear to be a universal feature, especially in smaller ADPKD cysts^4–7^. Importantly, many individuals with ADPKD continue to have residual PC1 expression because they carry missense (rather than inactivating) germline *PKD1* mutations^8,9,10^. As proof of principle, lowering the *Pkd1* dose is sufficient to produce PKD in mice, pigs, and monkeys^4,11–16^. Thus, if reduced dosage causes ADPKD, increasing the expression of the normal *PKD1* allele could arrest the disorder. However, despite this transformative potential, the factors governing *PKD1* dosage in ADPKD are mostly unknown, and currently, there are no mechanisms to activate the normal *PKD1* allele.

The 3’-untranslated region (3’-UTR), the mRNA portion that lies immediately downstream of the translation termination codon, protects the mRNA from degradation and facilitates translation through its poly(A) tail^17^. Paradoxically, the 3’-UTR can also mediate translation repression, chiefly *via* interaction with microRNAs (miRNAs) ^18–20^. Most mRNA 3’-UTRs harbor evolutionarily conserved miRNA-binding elements (MBEs), implying that cis-inhibition of translation is a pervasive mode of gene output regulation^21^. However, this intriguing aspect of the 3’-UTR function is poorly delineated. The prediction is that individual MBEs have a minor impact on host mRNA function, considering that miRNAs mostly act as rheostats and modestly repress mRNA targets. Counter to this prevailing logic, we reasoned that under certain circumstances, such as when gene dosage is already reduced due to haploinsufficiency, MBE-mediated cis-inhibition of the remaining allele could have a disease-modifying effect by governing the final protein output.

Our goal was to determine whether monoallelic *PKD1* derepression is possible and how it influences preclinical ADPKD progression. *PKD1* contains a miR-17 binding motif in its 3’-UTR, and miR-17 expression and activity are higher in ADPKD models^4,22–25^. Therefore, we tested the idea that *PKD1* mRNA is cis-inhibited by its 3’-UTR miR-17 motif, and blocking this autoinhibition will reverse the *PKD1* decline. We used CRISPR/Cas9 editing to delete this MBE from the *PKD1* gene in monoallelic ADPKD models. We found that eliminating the miR-17 motif is sufficient to improve *Pkd1* mRNA stability, raise Polycystin-1 (PC1) expression, and ameliorate cyst growth in cellular, *ex vivo*, and mouse PKD models. The other ADPKD gene, *PKD2,* also contains a 3’-UTR miR-17 binding motif; remarkably, deleting this MBE increases Polycystin-2 (PC2) levels and attenuates cyst growth in *Pkd1*-mutant models. Furthermore, acute pharmaceutical blockade of *Pkd1/2* cis-inhibition prevents cyst onset and stabilizes established PKD in mice. Finally, we demonstrate that *PKD1* or *PKD2* derepression reverses cyst-pathogenic events in patient-derived, primary ADPKD cell lines.

## RESULTS

### *Pkd1* is cis-repressed *via* its 3’-UTR miR-17 binding motif

The impact of individual 3’-UTR MBEs on host gene regulation is mostly unstudied. We explored the idea that cis-repression *via* its 3’-UTR miR-17 binding motif is a novel mechanism governing *Pkd1* dosage. We began by examining the impact of deleting this MBE in normal mouse kidneys. We designed sgRNAs that bind to *Pkd1* exon-46, flanking the DNA segment that encodes the miR-17 motif, and used CRISPR/Cas9 editing to generate *Pkd1* alleles (*Pkd1*^Δ17^) lacking the miR-17 binding site (**Fig. 1A**). First, we validated the motif deletion using DNA PCR followed by direct Sanger sequencing (**Fig. 1B-C**). Our editing approach did not inadvertently inactivate *Pkd1* since we observed normal kidney histology and renal function in 6-week-old and 18-week-old *Pkd1*^Δ17/Δ17^ mice (**Fig. 1D-F and Suppl. Fig. 1**). Despite the loss of the miR-17 binding site, PC1 expression was the same between kidneys of 6-week-old or 18-week-old *Pkd1*^Δ17/Δ17^ mice and their respective age-matched control *Pkd1*^+/+^ mice (**Suppl. Fig. 1D**), implying that there was no *Pkd1* cis-inhibition in the kidneys of adult mice.

**Figure 1:**
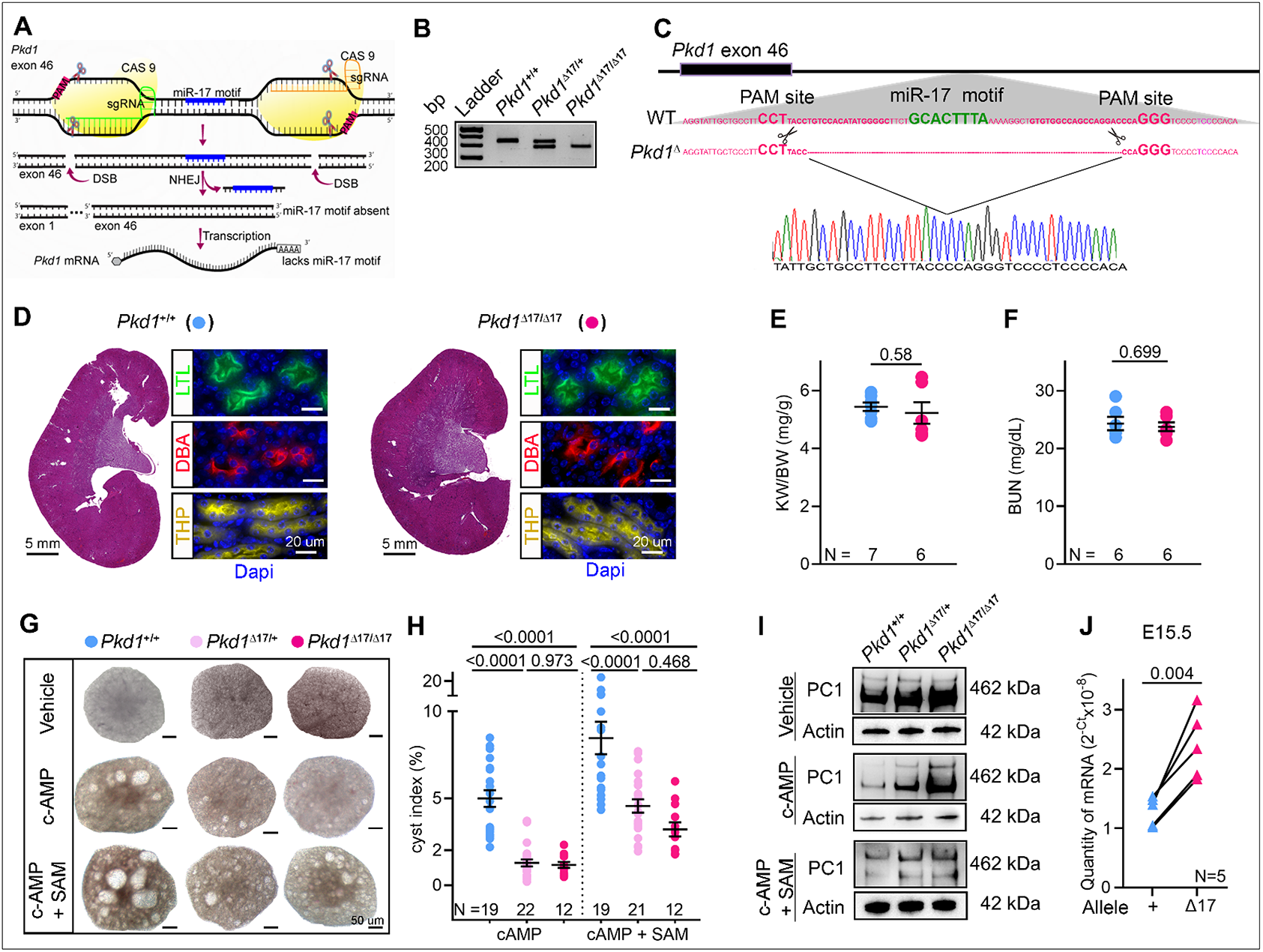
*Pkd1* mRNA is cis-repressed *via* its 3’-UTR miR-17 binding motif. (**A**) Graphic illustration of the CRISPR/Cas9 approach used to delete the miR-17 motif from *Pkd1* 3’-UTR (*Pkd1*^Δ17^). (**B**) PCR products obtained after amplification of tail DNA from mice with indicated genotypes. The lower band represents the Δ17 deletion. (**C**) 3’-UTR nucleotide sequence of wildtype (WT) and *Pkd1*^Δ17^ alleles. The miR-17 binding motif and sgRNA PAM sites are highlighted in bold green and pink, respectively. The pink dashed line indicates the deleted nucleotides in *Pkd1*^Δ17^. Sanger sequencing chromatogram depicting the nucleotide sequence of the *Pkd1*^Δ17^. (**D**) H&E staining, Lotus Tetragonolobus Lectin labeling (LTL, a proximal tubule marker), Tamm-Horsfall protein immunostaining (*THP*, a loop of Henle maker), and Dolichos Biflorus Agglutinin labeling (DBA, a collecting duct marker) showing normal kidney histology in 8-week-old *Pkd1*^+/+^ and *Pkd1*^Δ17/Δ17^ mice. (**E-F**) Normal kidney-weight-to-body-weight (KW/BW) and serum blood urea nitrogen (BUN) levels in 8-week-old *Pkd1*^+/+^ and *Pkd1*^Δ17/Δ17^ mice. (**G-H**) Images and cyst index quantification of E13.5 *Pkd1*^+/+^, *Pkd1*^Δ17/+^, and *Pkd1*^Δ17/Δ17^ kidneys grown for four days in culture media containing vehicle, 100 uM cAMP, or 100 uM cAMP plus 250 uM SAM. (**I**) Immunoblots depicting PC1 expression in *Pkd1*^+/+^, *Pkd1*^Δ17/+^, and *Pkd1*^Δ17/Δ17^ ex-vivo kidneys treated with vehicle, cAMP, or cAMP plus SAM. Actin is used as the loading control. (**J**) Allele-specific qRT-PCR showing the quantity of *Pkd1* mRNAs produced by the wildtype (+) and Δ17 alleles in E15.5 *Pkd1*^Δ17/+^ kidneys. Error bars indicate SEM. Statistical analysis: Student’s t-test (E and F); One-way ANOVA, Tukey’s multiple-comparisons test (H); Paired t-test (J).

miR-17 is abundantly expressed during embryogenesis, but its levels decline with postnatal kidney maturation. We reasoned that the lack of tonic *Pkd1* cis-inhibition was due to low basal miR-17 activity in mature kidneys. Therefore, we switched to analyzing embryonic (E) *Pkd1*^Δ17/+^ and *Pkd1*^Δ17/Δ17^ kidneys. We used *ex vivo* kidney organ culture to simultaneously assess the impact on *Pkd1* expression and cystogenesis. We cultured E13.5 littermate *Pkd1*^+/+^, *Pkd1*^Δ17/+^, and *Pkd1*^Δ17/Δ17^ kidneys for four days in media containing 100 μM 8-bromo-cAMP or 100 μM 8-bromo-cAMP plus S-adenosylmethionine

(SAM), or vehicle control (**Fig. 1G-H**). As expected ^26^, cAMP increased cyst formation in *Pkd1*^+/+^ kidneys compared to vehicle treatment, and this effect was further enhanced with the addition of SAM. Interestingly, the pro-cystogenic effect of cAMP and SAM was attenuated in *Pkd1*^Δ17/+^ and *Pkd1*^Δ17/Δ17^ kidneys (**Fig. 1G-H**). Moreover, we observed higher PC1 expression in *Pkd1*^Δ17/+^ and *Pkd1*^Δ17/Δ17^ compared to *Pkd1*^+/+^ *ex vivo* cultured kidneys by immunoblot analysis (**Fig. 1I**). These data indicate that miR-17 motif deletion derepresses PC1 and blocks the pro-cystogenic effect of cAMP in embryonic cultured kidneys.

Next, we measured the relative abundance of wild-type and Δ17 transcripts within the total *Pkd1* mRNA pool. We designed allele-specific primers taking advantage of the unique mRNA sequence created by CRISPR/Cas9 editing of the Δ17 allele. qRT-PCR revealed that in E15.5 heterozygous *Pkd1*^Δ17/+^ kidneys, the Δ17 allele contributed nearly 50% more transcripts than its wild-type counterpart (**Fig. 1J**). Similarly, we noted that the Δ17 allele produced more *Pkd1* mRNAs than the wild-type allele in *ex vivo Pkd1*^Δ17/+^ kidney cultures. This difference became even more pronounced in the presence of cAMP (**Suppl. Fig. 2A-D**). These observations further imply inhibition of wild-type *Pkd1* mRNAs but an evasion of repression and improved stability of *Pkd1*^Δ17^ mRNAs in embryonic kidneys.

### Endogenous monoallelic *Pkd1* derepression alleviates polycystic kidney disease

Kidney cyst formation ensues when *PKD1* dosage falls below a critical threshold. However, no approach exists to reverse the *PKD1* decline. Based on our observations in normal kidneys, we asked whether *Pkd1* is cis-repressed in ADPKD and if preventing this auto-inhibition has a disease-modifying impact. We first examined the *Pkd1*^RC/-^ cellular ADPKD model. This is a collecting duct-derived mouse cell line that harbors the missense RC mutation on one *Pkd1* allele, whereas the other allele is inactivated^27^. We used CRISPR/Cas9 editing to remove the 3’-UTR miR-17 motif from the RC allele (*Pkd1*^RCΔ17/-^) (**Suppl. Fig 3A-B**). We generated two independent *Pkd1*^RCΔ17/-^ clonal cell lines and characterized them both in relation to the unedited parental *Pkd1*^RC/-^ and *Pkd1*^RC/+^ cells. We previously reported that PC1 expression was reduced in *Pkd1*^RC/-^ cells compared to *Pkd1*^RC/+^ cells. Remarkably, eliminating the miR-17 motif restored PC1 expression in *Pkd1*^RCΔ17/-^ cell lines (**Fig. 2A** and **Suppl. Fig 4A**). Next, employing several independent assays, we demonstrated that this degree of PC1 derepression was sufficient to reverse pathogenic sequelae of PKD. First, we noted a higher proliferation rate and 3D cyst size in *Pkd1*^RC/-^ cells than in *Pkd1*^RC/+^ cells, which was normalized after PC1 derepression in *Pkd1*^RCΔ17/-^ cells (**Fig. 2B-C and Suppl. Fig 4B-D**). Second, we observed that while cAMP, glucose, and SAM increased the already elevated proliferation of *Pkd1*^RC/-^ cells, *Pkd1*^RCΔ17/-^ cells were resistant to these pro-proliferative stimuli (**Fig. 2D and Suppl. Table 1**). Third, we used MitoTracker to assess mitochondrial membrane potential as a proxy of oxidative phosphorylation and anti-pCreb1 antibody immunofluorescence as a readout of c-AMP signaling. Compared to *Pkd1*^RC/+^ cells, we observed a reduced MitoTracker signal and higher pCreb1 expression in *Pkd1*^RC/-^ cells. The opposite was true for *Pkd1*^RCΔ17/-^ cells, which exhibited restored MitoTracker signal and lowered pCreb1 expression (**Fig. 2E** and **Suppl. Fig 4E**). Finally, immunoblot analysis revealed elevated Yap1, pCreb1, and c-Myc expression in *Pkd1*^RC/-^ cells compared to *Pkd1*^RC/+^ cells, which returned to baseline in *Pkd1*^RCΔ17/-^ cells (**Suppl. Fig 4F**).

**Figure 2:**
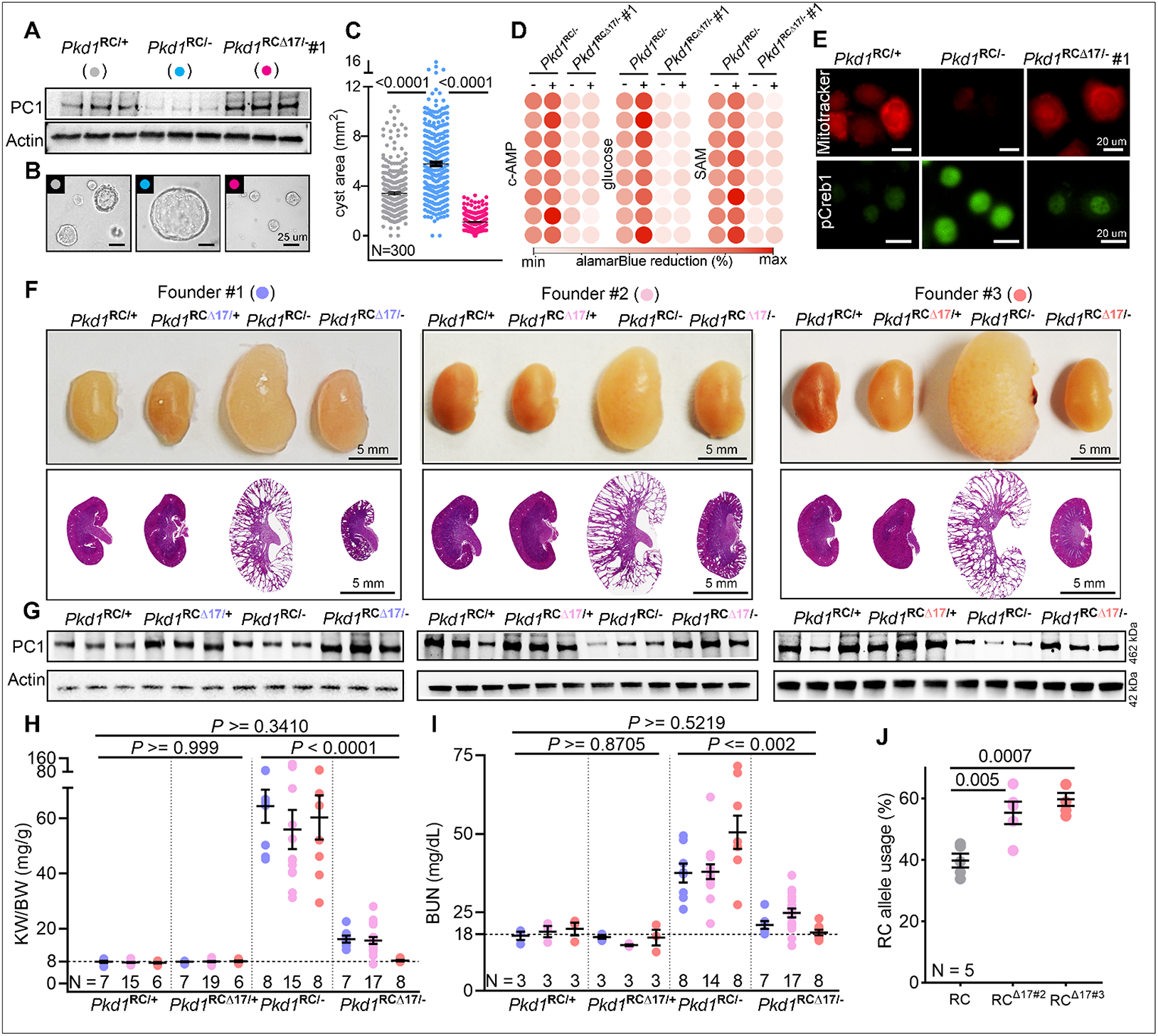
Monoallelic *Pkd1* derepression alleviates polycystic kidney disease: (**A**) Immunoblot showing reduced PC1 expression in *Pkd1*^RC/-^ compared to *Pkd1*^RC/+^ cells. PC1 level was restored in *Pkd1*^RCΔ17/-^ cells. Actin serves as the loading control. (**B-C**) Representative images and quantification showing increased 3D cyst size of *Pkd1*^RC/-^ compared to *Pkd1*^RC/+^ cells cultured in Matrigel. Cyst size was normalized in *Pkd1*^RCΔ17/-^ cells. (**D**) Heatmap showing alamarBlue-assessed proliferation of *Pkd1*^RC/-^ and *Pkd1*^RCΔ17/-^ cells in the absence (−) or presence (+) of 100 uM cAMP, 17 mM glucose, or 100 uM SAM. N=8, each circle represents a biological replicate. (**E**) Representative images showing Mito tracker labeling and anti-pCreb1 immunostaining in *Pkd1*^RC/+^, *Pkd1*^RC/-^, and *Pkd1*^RCΔ17/-^ cells. (**F**) Gross kidney and H&E-stained kidney sections from 18-day-old mice with the indicated genotypes. Data from the progeny of the three founders are shown separately. (**G**) Immunoblot showing PC1 expression in kidneys of 18-day-old mice with the indicated genotypes derived from the three founders. Actin serves as the loading control. (**H-I**) KW/BW ratio and BUN levels in mice with the indicated genotypes are shown. Data from all three founders are shown. Founder#1 (blue circles), Founder#2 (light pink circles), and Founder#3 (orange circles). (**J**) Paired-end RNA-seq data showing the RC allele usage in *Pkd1*^RC/-^ (grey circles), *Pkd1*^RCΔ17/-^ founder#2 (pink circles), and *Pkd1*^RCΔ17/-^ founder#3 (orange circles). Error bars indicate SEM. Statistical analysis: One-way ANOVA, Tukey’s multiple-comparisons test (C, H, I, J).

**Figure 3:**
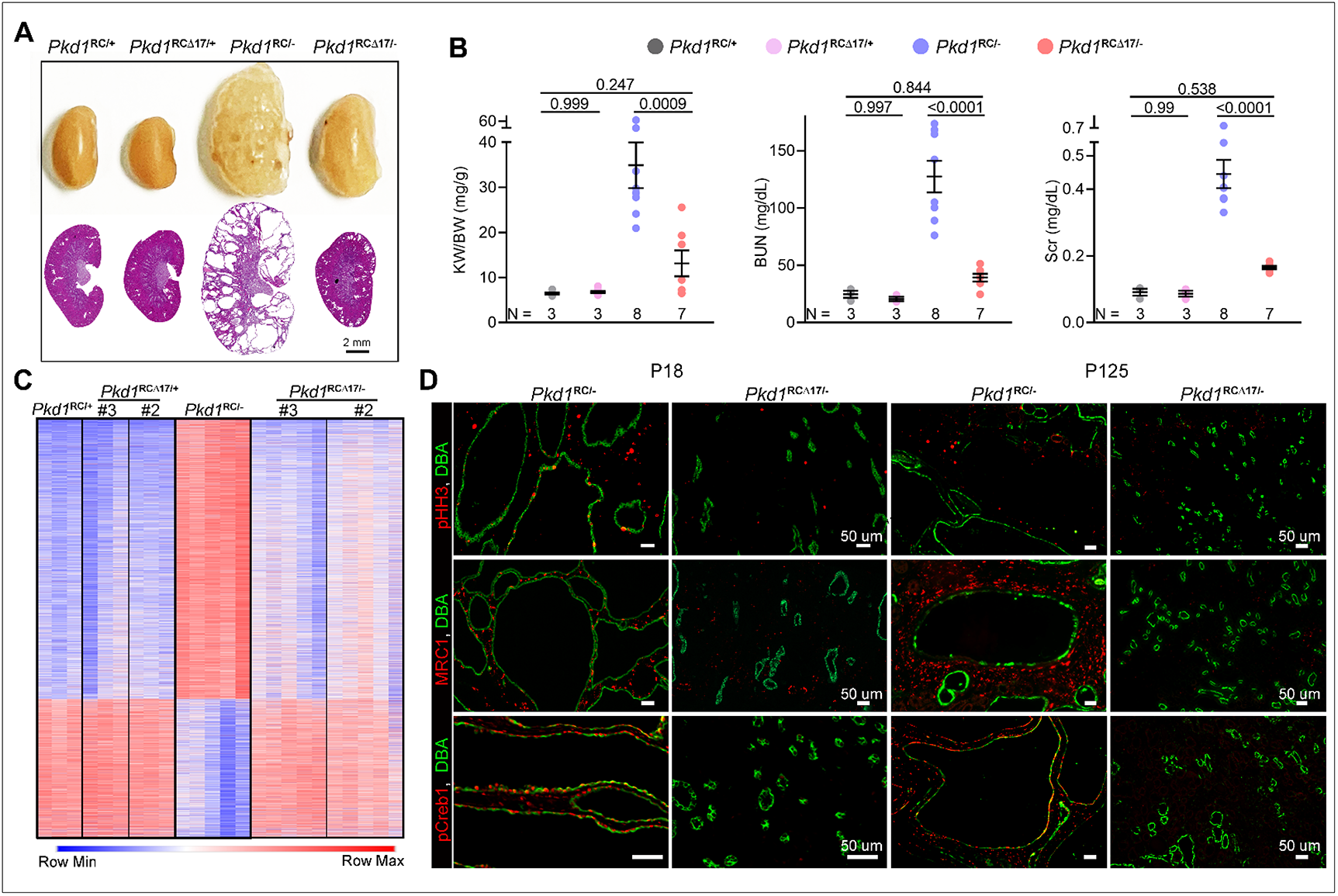
*Pkd1* derepression attenuates disease progression and reverses cyst-pathogenic events: (**A**) Gross kidney images and H&E-stained kidney sections from 18-week-old mice with the indicated genotypes derived from founder#3. miR-17 motif deletion was associated with sustained benefit and suppressed long-term PKD progression. (**B**) KW/BW, BUN, and serum creatinine (Scr) levels in the 18-week-old progeny of founder#3. (**C**) Heatmap showing global mRNA expression profiles of kidneys from 18-day-old mice with the indicated genotypes. #3 = founder#3; #2 = founder#2. mRNAs that were dysregulated in *Pkd1*^RC/-^ compared to *Pkd1*^RC/+^ kidneys but exhibited improved expression in *Pkd1*^RCΔ17/-^ kidneys were chosen for visualization. (**D**) Representative images showing phospho-Histone-H3 (pHH3), Mannose Receptor C-Type 1 (MRC1), or pCreb1 immunostaining in kidney sections of 18-day-old and 18-week-old *Pkd1*^RC/-^ and *Pkd1*^RCΔ17/-^ mice. The sections were co-labeled with DBA to mark collecting duct-derived cysts. Error bars indicate SEM. Statistical analysis: One-way ANOVA, Tukey’s multiple-comparisons test (B).

**Figure 4:**
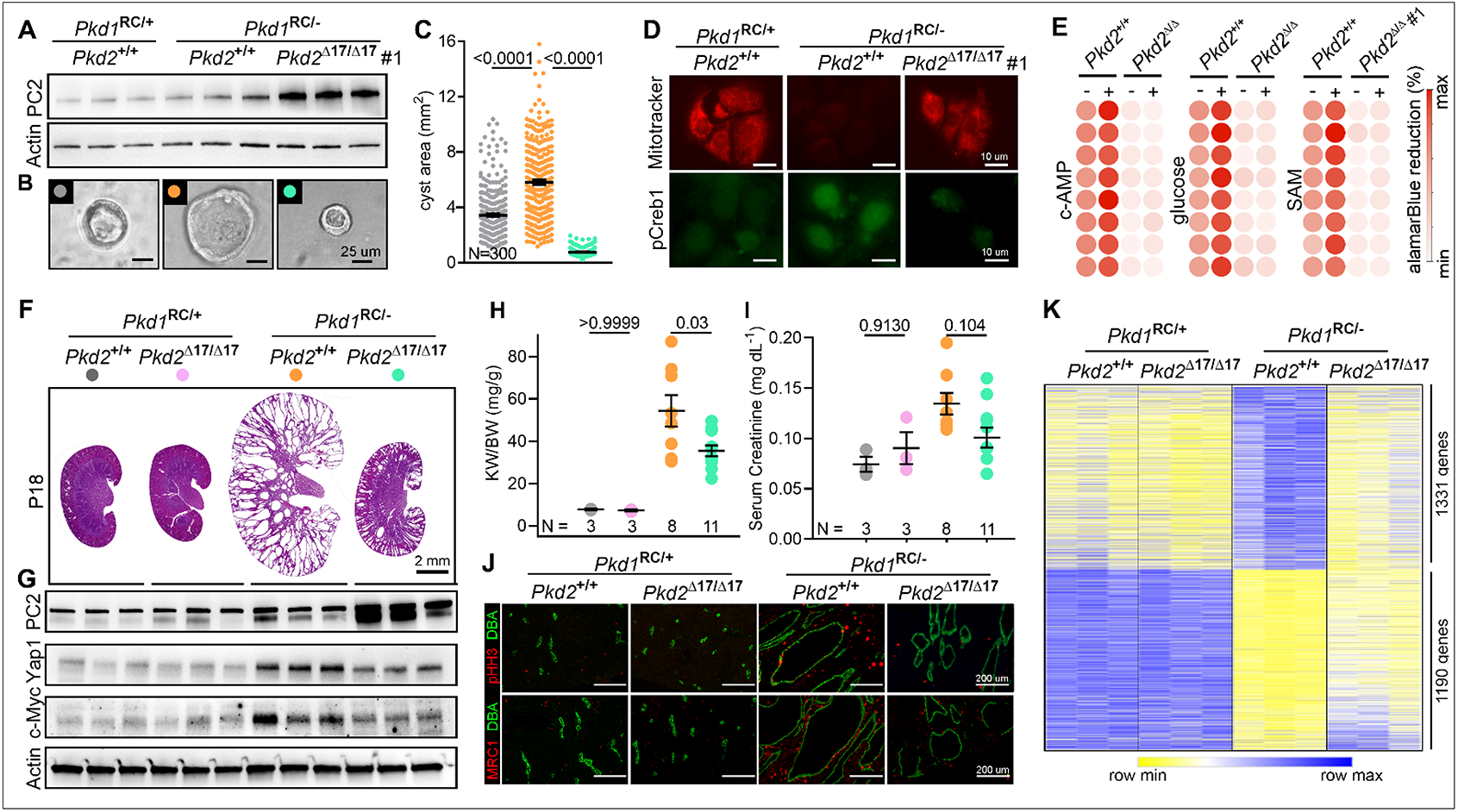
*Pkd2* derepression attenuates cyst growth in *Pkd1*-mutant models: (**A**) Immunoblot showing Polycystin-2 (PC2) expression in cells with the indicated genotypes. miR-17 motif deletion from *Pkd2* 3’-UTR leads to higher PC2 expression in *Pkd1*^RC/-^ cells. Actin is used as the loading control. (**B-C**) Representative images and quantification showing 3D cyst size of cells with indicated genotypes grown in Matrigel cultures. (**D**) Representative images showing mitotracker labeling and anti-pCreb1 immunostaining in cells with the indicated genotypes. (**E**) Heatmap showing alamarBlue-assessed proliferation of *Pkd1*^RC/-^ and *Pkd1*^RC/-^; *Pkd2*^Δ17/Δ17^ cells in the absence (−) or presence (+) of cAMP, glucose, or SAM. N=8, each circle represents a biological replicate. (**F**) H&E-stained kidney sections from 18-day-old mice with the indicated genotypes are shown. (**G**) Immunoblots showing PC2, Yap1, and c-Myc expression in kidneys of 18-day-old mice with the indicated genotypes. (**H-I**) KW/BW and serum creatinine levels in 18-day-old mice with the indicated genotypes. (**J**) Representative images showing pHH3 and MRC1 immunostaining in kidney sections from 18-day-old mice with the indicated genotypes. (**K**) Heatmap showing differential mRNA expression in kidneys of 18-day-old mice with the indicated genotypes. mRNAs that were dysregulated in *Pkd1*^RC/-^ compared to *Pkd1*^RC/+^ kidneys but exhibited improved expression in *Pkd1*^RC/-^*; Pkd2*^Δ17/Δ17^ kidney were chosen for heatmap visualization. Error bars indicate SEM. Statistical analysis: One-way ANOVA, Tukey’s multiple-comparisons test (B, H, and I).

Based on these encouraging results, we next modeled the 3’-UTR Δ17 deletions *in vivo*. We CRISPR-edited Ksp^Cre+^; *Pkd1*^RC/RC^ fertilized eggs to eliminate the miR-17 motif from the *Pkd1* RC allele. We implanted these eggs into pseudopregnant surrogate female mice and eventually obtained three germline-transmitting heterozygous Ksp^Cre+^; *Pkd1*^RCΔ17/RC^ founder mice. Using DNA PCR and Sanger sequencing, we validated that the miR-17 motif was indeed deleted in all three founder mice from one RC allele, whereas the other RC allele still contained the wild-type 3’-UTR (**Suppl. Fig 5**). We selected mice with heterozygous Δ17 deletions because breeding them with *Pkd1*^F/F^ mice allowed us to generate the following four relevant genotypes from the same breeding pair: *Pkd1*^RC/F^ (*Pkd1*^RC/+^), *Pkd1*^RCΔ17/F^ (*Pkd1*^RCΔ17/+^), Ksp^Cre+^; *Pkd1*^RC/F^ (*Pkd1*^RC/-^), and Ksp^Cre+^; *Pkd1*^RCΔ17/F^ (*Pkd1*^RCΔ17/-^). Data from the 18-day-old progeny of all three founders are shown in **Fig. 2F-H**. First, we noted normal kidney histology and function in *Pkd1*^RCΔ17/+^ mice, again indicating that miR-17 motif deletion does not disrupt *Pkd1* or produce PKD (**Fig. 2F**). For each founder progeny, we observed reduced PC1 expression, severe cystic kidney disease, an increased kidney-weight-to-body-weight (KW/BW) ratio, and higher serum BUN levels in *Pkd1*^RC/-^ mice than in *Pkd1*^RC/+^ mice (**Fig. 2F-J**). As with the cell lines, miR-17 motif deletion caused PC1 derepression in *Pkd1*^RCΔ17/-^ kidneys compared to *Pkd1*^RC/-^ kidneys (**Fig. 2G**). Moreover, using paired-end RNA-seq, we noted higher RC allele usage in *Pkd1*^RCΔ17/-^ kidneys than in *Pkd1*^RC/-^ kidneys, further indicating *Pkd1* derepression (**Fig. 2J**). Strikingly, the cystic disease was almost completely alleviated, and KW/BW and serum BUN were nearly normalized in *Pkd1*^RCΔ17/-mice^ compared to *Pkd1*^RC/-^ mice (**Fig. 2F-I**). To examine the long-term effects, we prospectively followed founder #2 and #3 progeny for 8 and 18 weeks, respectively. Founder#2 progeny exhibited an aggressive cystic disease phenotype, with 76.4% (13/17) of *Pkd1*^RC/-^ mice succumbing to kidney failure before 8 weeks of age (**Suppl. Fig. 6A-B**). Moreover, the four surviving *Pkd1*^RC/-^ mice exhibited severe PKD and near-fatal kidney failure. In contrast, only 27.2% (6/22) of *Pkd1*^RCΔ17/-^ mice died by 8 weeks, and the surviving mice had fewer cysts and relatively preserved kidney function (**Suppl. Fig. 6C-E**). All founder#3 *Pkd1*^RC/-^ progeny survived until 18 weeks of age. However, they developed progressive kidney failure, as evidenced by an average blood urea nitrogen (BUN) of >100 mg/dl and serum creatinine of >0.4 mg/dl (**Fig 3B**). Founder #3 *Pkd1*^RCΔ17/-^ mice exhibited minimal disease progression with average BUN <30 mg/dl and serum creatinine <0.2 mg/dl (**Fig. 3A-B**).

Large-scale transcriptomic dysregulation, activation of tubular proliferation and oncogenic signaling, and interstitial inflammation are some of the key pathological hallmarks of ADPKD. We next addressed whether these changes were blunted by PC1 derepression. We performed RNA-seq analysis using kidney samples from 18-day-old *Pkd1*^RC/+^, *Pkd1*^RCΔ17/+^, *Pkd1*^RC/-^, and *Pkd1*^RCΔ17/-^ mice. We observed dysregulation of an extensive network of gene transcripts with upregulation of 4157 and downregulation of 2067 mRNAs in cystic *Pkd1*^RC/-^ compared to noncystic *Pkd1*^RC/+^ control kidneys (**Fig. 3C**). Mirroring kidney histology, *Pkd1*^RCΔ17/+^ exhibited a nearly identical gene expression pattern as *Pkd1*^RC/+^ kidney. Impressively, >95% of dysregulated mRNAs in *Pkd1*^RC/-^ kidneys showed improved (or normalized) expression in *Pkd1*^RCΔ17/-^ kidneys (**Fig. 3C**). Consistent with the RNA-seq data, immunoblot analysis revealed reduced c-Myc and Yap1 in the kidneys of 18-day-old *Pkd1*^RCΔ17/-^ mice compared to *Pkd1*^RC/-^ mice (**Suppl. Fig. 6F**). Finally, immunofluorescence analysis demonstrated fewer anti-phospho-Histone-H3-positive cells, indicating lower proliferation, and reduced anti-pCreb1 and anti-MRC1 signals, implying attenuated c-AMP signaling and cyst-associated inflammation, respectively, in kidneys of 18-day-old and 18-week-old *Pkd1*^RCΔ17/-^ mice compared to *Pkd1*^RC/-^ mice (**Fig. 3D**).

### Preventing *Pkd2* cis-inhibition attenuates cyst growth in *Pkd1*-mutant models

A long-standing question has been whether increasing PC2 can compensate for low PC1. Interestingly, similar to *PKD1*, *PKD2* harbors an evolutionarily conserved 3’-UTR miR-17 motif. Our work indicates heightened miR-17 repressive activity in *Pkd1*-mutant ADPKD models. Therefore, we wondered whether *Pkd2* is cis-inhibited and whether preventing this autoinhibition positively impacts disease progression in *Pkd1*-mutant models. To address these questions, we again began with the *Pkd1^RC/-^* cellular model but this time used CRISPR/Cas9 to delete the miR-17 motifs from the *Pkd2* 3’-UTR (*Pkd1*^RC/-^; *Pkd2*^Δ17/Δ17^) while keeping the *Pkd1* miR-17 motif intact (**Suppl. Fig 7A-B**). Consistent with 3’-UTR cis-inhibition, using qRT-PCR and immunoblot analysis, we observed higher *Pkd2* and PC2 expression in two *Pkd1*^RC/-^; *Pkd2*^Δ17/Δ17^ clonal cell lines compared to their unedited parental *Pkd1*^*RC/-*^ cells (**Fig. 4A** and **Suppl. Fig. 8A-B**). PC1 expression remained unchanged between the edited and unedited cells, indicating the specificity of miR-17 motif deletion from *Pkd2* 3’-UTRs (**Suppl. Fig. 8E**). Surprisingly, we noted that PC2 derepression was associated with reduced 3D cyst growth, restored MitoTracker signal, and downregulation of pCreb1, Yap1, Mettl3, and c-Myc expression in *Pkd1*^RC/-^; *Pkd2*^Δ17/Δ17^ cells compared to *Pkd1*^RC/-^ cells (**Fig. 4B-D** and **Suppl. Fig. 8C-F**). As was the case with the *Pkd1* 3’-UTR deletions, while cAMP, glucose, and SAM promoted proliferation of *Pkd1^RC/-^* cells, we noted that this stimulatory effect was lost in *Pkd1^RC/-^*; *Pkd2*^Δ17/Δ17^ cells (**Fig. 4E, Suppl. Table 2**).

Our observations in *Pkd1*^RC/-^ cells suggest a tantalizing possibility that preventing *Pkd2* cis-inhibition and improving PC2 expression compensates and retards disease progression in *Pkd1*-mutant models. We tested this notion *in vivo* by deleting the *Pkd2* 3’-UTR miR-17 motif (*Pkd2*^Δ17/Δ17^) in *Pkd1*^RC/-^ mice. Briefly, we CRISPR/Cas9-edited Ksp^Cre^; *Pkd1*^RC/RC^ mice to generate Ksp^Cre^; *Pkd1*^RC/RC^; *Pkd2*^Δ17/+^ mice (see **Suppl. Fig. 9** for details). These mice were then bred with *Pkd1*^F/F^ mice to eventually generate the following four genotypes: (i) *Pkd1*^RC/F^; *Pkd2*^+/+^, (ii) *Pkd1*^RC/F^; *Pkd2*^Δ17/Δ17^, (iii) Ksp^Cre^; *Pkd1*^RC/F^; *Pkd2*^+/+^, and (iv) Ksp^Cre^; *Pkd1*^RC/F^; *Pkd2*^Δ17/Δ17^. Characterization of these mice revealed that *Pkd2* miR-17 motif deletion in the noncystic setting did not cause PC2 upregulation, and both *Pkd1*^RC/F^; *Pkd2*^+/+^ and *Pkd1*^RC/F^; *Pkd2*^Δ17/Δ17^ mice exhibited normal kidney histology and function (**Fig. 4F and G**). In contrast, *Pkd2* miR-17 motif deletion in cystic *Pkd1*^RC/-^ mice was associated with higher PC2 expression. Moreover, KW/BW and serum creatinine levels were reduced by 34.8% and 25%, respectively, in *Pkd1*^RC/-^; *Pkd2*^Δ17/Δ17^ compared to *Pkd1*^RC/-^; *Pkd2*^+/+^ mice (**Fig. 4H-I**). Consistently, we observed that compared to *Pkd1*^RC/-^; *Pkd2*^+/+^ kidneys, *Pkd1*^RC/-^; *Pkd2*^Δ17/Δ17^ kidneys exhibited reduced c-Myc and Yap1 expression (**Fig. 4G**) and lower cyst proliferation and interstitial inflammation (**Fig. 4J**). As an additional phenotypic characterization, we performed RNA-seq analysis to compare the kidney transcriptomic profile in the four groups of mice (**Fig. 4K**). The mRNA expression patterns were nearly identical in *Pkd1*^RC/F^; *Pkd2*^+/+^ and *Pkd1*^RC/F^; *Pkd2*^Δ17/Δ17^, further implying that *Pkd2* miR-17 motif elimination has minimal impact in noncystic kidneys. As expected, the cystic *Pkd1*^RC/-^; *Pkd2*^+/+^ kidneys exhibited widespread mRNA dysregulation compared to noncystic *Pkd1*^RC/+^; *Pkd2*^+/+^ control kidneys. We found that *Pkd2* miR-17 motif deletion was associated with improved expression of nearly 50% of these dysregulated mRNAs in *Pkd1*^RC/-^; *Pkd2*^Δ17/Δ17^ (**Fig. 4K**).

### Acute blockade of *Pkd1* and *Pkd2* cis-inhibition ameliorates PKD

Our CRISPR-edited clonal cellular or mouse ADPKD models lead to chronic *Pkd1* or *Pkd2* derepression. Therefore, we could not assess whether acute derepression of *Pkd1*/*2,* just as the cysts are forming, will prevent disease onset or if restoring *Pkd1*/*2* can reign in established PKD. To answer these questions, we employed the anti-miR-17 oligonucleotide RGLS4326 as a tool to acutely block *Pkd1* and *Pkd2* cis-inhibition^28^. First, we confirmed that compared to vehicle (PBS) or control oligonucleotide, RGLS4326 indeed increased *Pkd1/2* and PC1/2 expression in *Pkd1*^RC/-^ cells (**Fig 5A-B**). The *Pkd1/2-*boosting effect of RGLS4326 was apparent within three days after treatment. Moreover, RGLS4326-treated *Pkd1*^RC/-^ cells had reduced proliferation and produced smaller cysts in 3D Matrigel cultures than PBS- or control oligonucleotide-treated *Pkd1*^RC/-^ cells (**Suppl. Fig 10A-C**). The cyst-reducing effect of RGLS4326 was present but blunted in *Pkd1*^RCΔ17/-^ or *Pkd1*^RC/-^; *Pkd2*^Δ17/Δ17^ cell lines, suggesting that this compound mediates its benefits in *Pkd1*^RC/-^ cells primarily via *Pkd1/2* derepression (**Suppl. Fig 10D-G**). Next, we tested the impact of acutely raising *Pkd1/2* in *Pkd1*^RC/-^ cells after cysts had already formed. We cultured untreated *Pkd1*^RC/-^ cells in Matrigel for four days, allowing the cyst to grow. We then treated these cysts with a vehicle, control oligonucleotide, or RGLS4326 and monitored them for three additional days. Vehicle and control oligonucleotide-treated cysts nearly tripled in size, whereas RGLS4326 treatment suppressed this growth (**Fig 5C**). RGLS4326 treatment was also associated with lower Yap1, c-Myc, and pCreb1 expression and a higher mitotracker signal, indicating that acute *Pkd1/2* derepression reverts cyst-pathogenic events (**Suppl. Fig 10H-I**).

**Figure 5:**
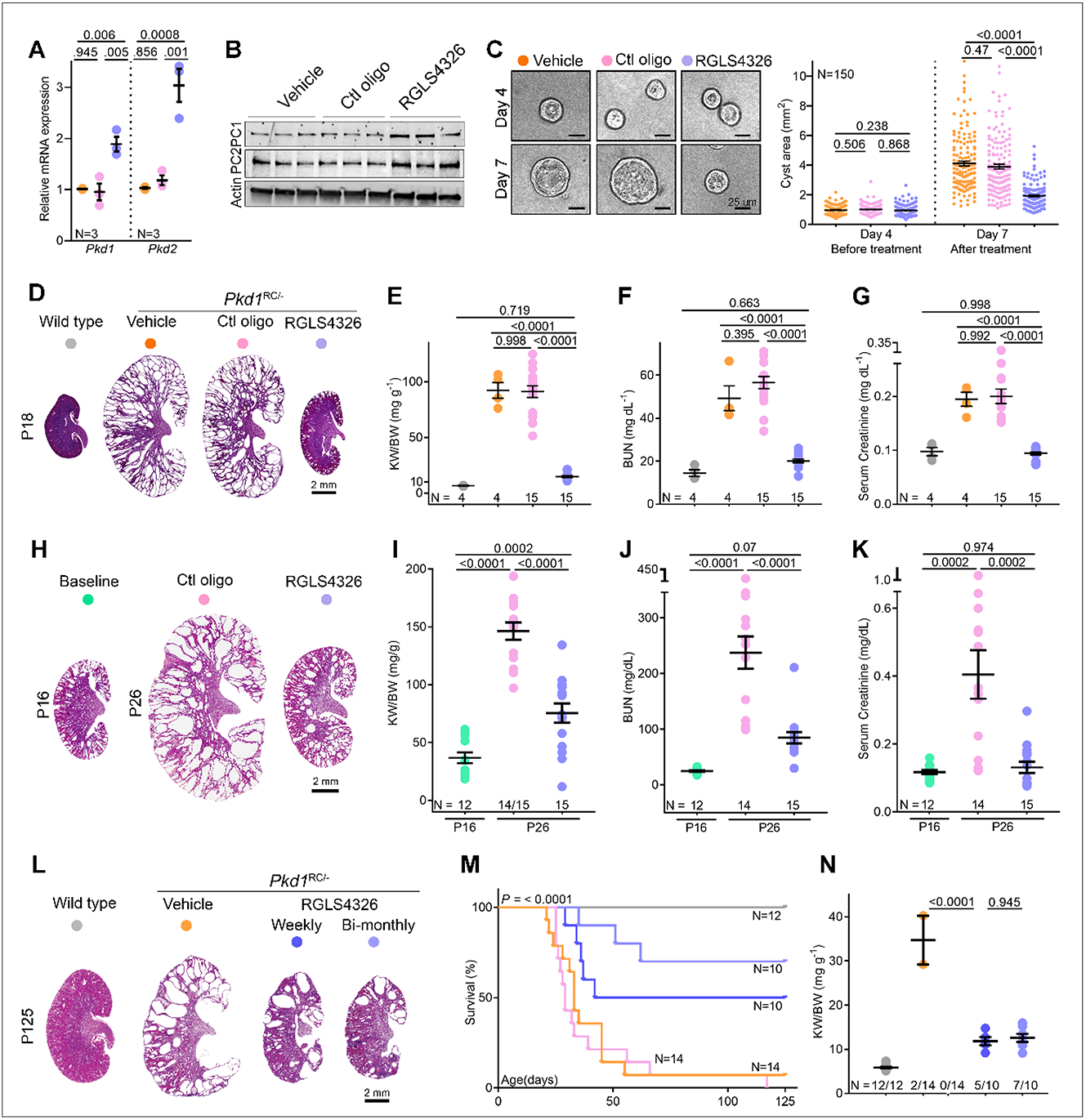
Acute *Pkd1* and *Pkd2* derepression attenuate cyst growth: (**A-B**) qRT-PCR and immunoblot analysis showing *Pkd1*/PC1 and *Pkd2*/PC2 expression in *Pkd1*^RC/-^ cells transfected with vehicle (PBS), 100 uM control oligonucleotide, or 100 uM RGLS4326. (**C**) Representative images and quantification of 3D cyst size of *Pkd1*^RC/-^ cells cultured in Matrigel before (day4) or after (day7) transfection with vehicle (PBS), 100 uM control oligonucleotide, or 100 uM RGLS4326. (**D**) H&E-stained kidney sections from 18-day-old *Pkd1*^RC/-^ mice injected on P10, P11, P12, and P16 either with vehicle (PBS), 20 mg/kg control oligonucleotide, or 20 mg/kg RGLS4326. H&E-stained kidney section from untreated 18-day-old wildtype mouse is shown for reference. (**E-G**) KW/BW, BUN, and serum creatinine levels in 18-day-old *Pkd1*^RC/-^ mice treated with vehicle (PBS), 20 mg/kg control oligonucleotide, or 20 mg/kg RGLS4326 are shown. Data from untreated 18-day-old wildtype mice are shown as a reference. (**H**) H&E-stained kidney sections from 26-day-old *Pkd1*^RC/-^ mice injected on P16 and P17 with 20 mg/kg control oligonucleotide or 20 mg/kg RGLS4326. H&E-stained kidney sections from genetically matched but untreated 16-day-old *Pkd1*^RC/-^ mice are shown to depict disease prior to starting treatment. (**I-K**) KW/BW, BUN, and serum creatinine levels in untreated 16-day-old or treated 26-day-old *Pkd1*^RC/-^ mice are shown. (**L-N**) *Pkd1*^RC/-^ mice were injected on P16 and P17 with vehicle, 20 mg/kg RGLS4326 or 20 mg/kg control oligonucleotide. These mice then received their respective treatment regimen every week until 18-weeks of age. The fourth cohort of *Pkd1*^RC/-^ mice received 20 mg/kg RGLS4326 treatment on P16, P17, and bimonthly thereafter. (**L**) H&E-stained kidney sections of 125-day-old *Pkd1*^RC/-^ mice on the vehicle or RGLS4326 treatment are shown. (**M**) Kaplan-Meir survival curves of *Pkd1*^RC/-^ mice in the four treatment groups are shown. Survival of untreated wildtype is shown as a reference. (**N**) KW/BW ratio in mice that survived till 125 days is shown. Error bars indicate SEM. Statistical analysis: One-way ANOVA, Tukey’s multiple-comparisons test (A, C, E-G, I-K, and N); Mantel-Cox (M).

We next examined whether the observations in cells can be replicated *in vivo*. In the first study, we treated *Pkd1*^RC/-^ mice with vehicle (PBS), control oligonucleotide, or RGLS4326 starting at P10, the age at which cysts begin to form in this model. By P18, we noted marked kidney enlargement with a >10-fold higher KW/BW ratio and elevated BUN and serum creatinine in PBS- and control oligonucleotide-treated mice compared to age-matched wild-type mice (**Fig. 5D-G**). Strikingly, PKD was virtually prevented, and renal function remained normal in P18 RGLS4326-treated *Pkd1*^RC/-^ mice (**Fig. 5D-G**). In the second study, we began treatment at P16 when *Pkd1*^RC/-^ mice had already developed cystic disease. By P26, 1/15 control oligonucleotide-treated mice had died, and the surviving mice had developed progressive kidney enlargement and near-fatal kidney failure. In contrast, we observed attenuation of PKD progression and stabilization of kidney function in RGLS4326-treated *Pkd1*^RC/*-*^-KO mice (**Fig. 5H-K**). Finally, in a third study, we assessed the long-term effects of *Pkd1/2* derepression in mice that had already developed PKD. We treated *Pkd1*^RC/-^ mice on P16 and P17 with vehicle, 20 mg/kg RGLS4326 or 20 mg/kg control oligonucleotide. These mice then received their respective treatment regimens every week until 18 weeks of age. A fourth group of *Pkd1*^RC/-^ mice received 20 mg/kg RGLS4326 treatment on P16 and P17 and every other week thereafter. A total of 85.7% (12/14) of PBS-treated and 100% (14/14) of control oligonucleotide-treated *Pkd1*^RC/-^-KO mice succumbed to their disease before 18 weeks of age. In contrast, 70% (7/10) and 50% (5/10) of *Pkd1*^RC/-^ mice treated with RGLS4326 bi-monthly or weekly, respectively, survived until 18 weeks of age (**Fig. 5M**). Furthermore, we noted substantially preserved kidney parenchyma (**Fig. 5L**) and reduced KW/BW (**Fig. 5N**) among the surviving mice in the RGLS4326 group. Thus, acute pharmaceutical *Pkd1/2* derepression, including after cyst onset, attenuates murine PKD.

### *PKD1*Δ_17_ or *PKD2*Δ_17_ alleles reduce cyst growth of patient-derived primary ADPKD cultures

A logical but crucial question is whether *PKD1/2* cis-inhibition is a feature of human ADPKD. To assess the translational potential of our findings in mice, we turned to primary human ADPKD cultures. We derived these cells from cysts of freshly discarded ADPKD nephrectomy samples from four affected individuals. First, we designed human-specific sgRNAs targeting the miR-17 motifs in the *PKD1* or *PKD2* 3’-UTRs. We then used CRISPR/Cas9 editing to eliminate the *PKD1* or *PKD2* miR-17 motif in primary ADPKD cultures from all four donors (**Suppl. Fig 11**). We noted higher PC1 levels within three days of modeling the *PKD1*^Δ17^ allele in all four CRISPR-edited cultures compared to their respective unedited parental controls (**Fig. 6A**). Similarly, modeling *PKD2*^Δ17^ alleles led to higher PC2 expression in edited versus unedited ADPKD cultures (**Fig. 6B**). We next assessed the functional significance of *PKD1* or *PKD2* derepression by performing Matrigel 3D cystogenesis, alamarBlue proliferation assays, live-cell MitoTracker labeling, and anti-pCREB1 immunofluorescence. The *PKD1*^Δ17^ or *PKD2*^Δ17^ cells formed smaller cysts (**Fig. 6C-F**) and exhibited lower proliferation rates (**Suppl. Fig. 12**), higher MitoTracker signal, and lower pCREB1 expression compared to their respective unedited controls (**Fig. 6G-H**). These data imply ongoing *PKD1/2* cis-inhibition and the potential benefit of derepressing *PKD1/2* in human ADPKD cells.

**Figure 6:**
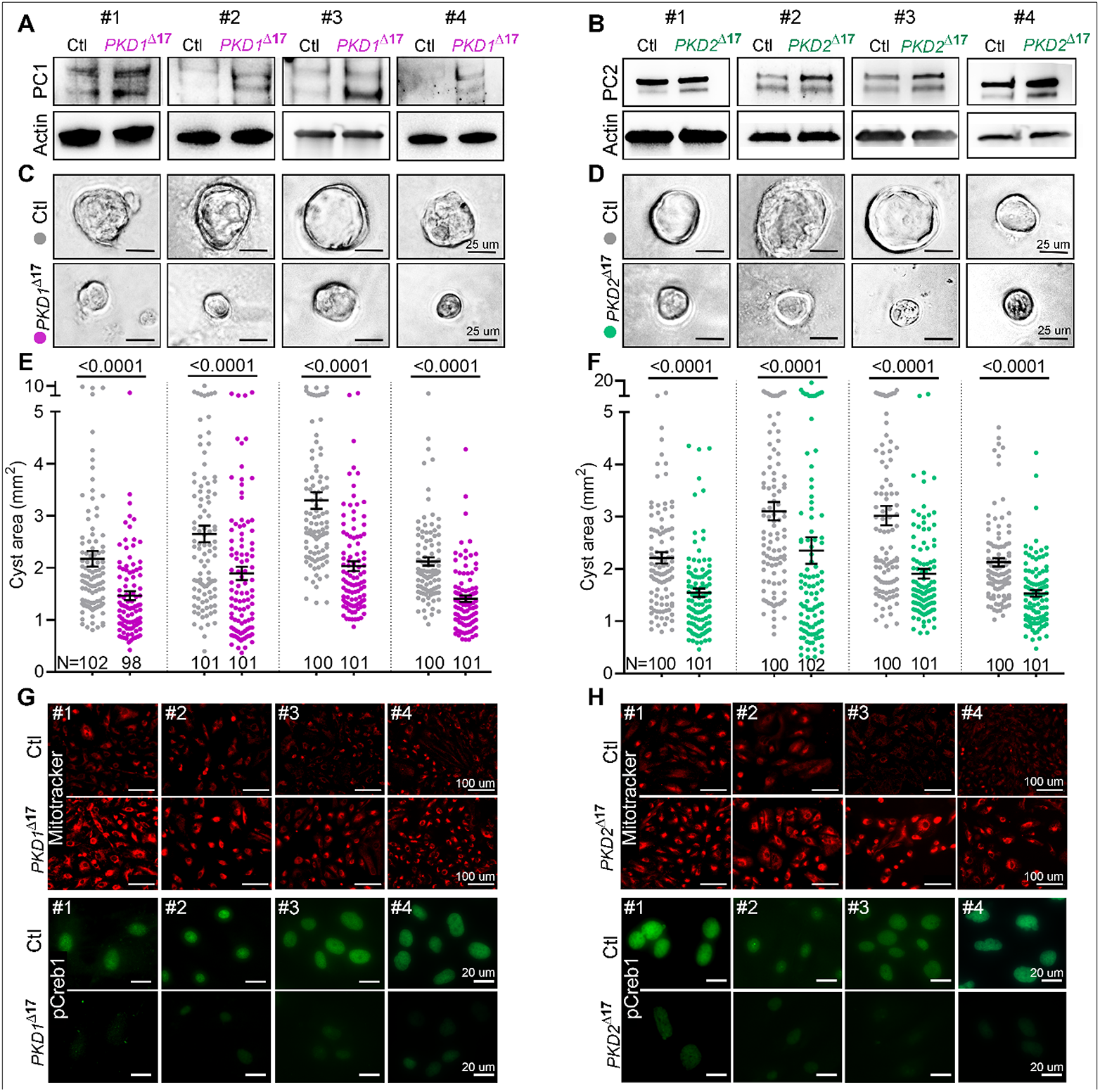
*PKD1*^Δ17^ or *PKD2*^Δ17^ revert cyst-pathogenic sequala in primary human ADPKD cultures. CRISPR/Cas9-editing was used to delete the miR-17 motif from *PKD1* 3’-UTR (*PKD1*^Δ17^) or *PKD2* 3’-UTR (*PKD2*^Δ17^) in primary ADPKD cultures from four human donors (#1 through #4). (**A-B**) Immunoblots showing higher PC1 expression in *PKD1*^Δ17^ and higher PC2 expression in *PKD2*^Δ17^ ADPKD cultures compared to their respective unedited parental ADPKD cultures. Protein bands are at 460 kDa (**A**) and 110-120 kDa size (**B**). Actin serves as the loading control. (**C-F**) Images and quantification showing reduced cyst size of *PKD1*^Δ17^ and *PKD2*^Δ17^ compared to their respective unedited parental ADPKD cultures. (**G-H**) Images showing higher mitotracker labeling (red) and reduced pCREB1 immunostaining (green) in *PKD1*^Δ17^ and *PKD2*^Δ17^ ADPKD cultures compared to their respective unedited parental ADPKD cultures. Errors bars represent SEM, Statistical analysis: Students *t*-test (E-F).

## DISCUSSION

*PKD1* loss-of-function as the genetic cause of ADPKD was discovered over 25 years ago^29–31^, but approaches to restore *PKD1* expression remain elusive. We provide a feasible framework for increasing endogenous *PKD1* and show for the first time that monoallelic *Pkd1* derepression is sufficient to alleviate preclinical PKD. As an attractive safety feature, *Pkd1* cis-inhibition appears to be an ADPKD-specific phenomenon since we observed that the *Pkd1* miR-17 motif has minimal impact in wild-type adult mouse kidneys. Furthermore, our work suggests that restoring even hypomorphic *Pkd1* mutants may be beneficial. On a cautionary note, particularly for modalities employing exogenous *PKD1* supplementation, raising *Pkd1* above wild-type levels produces cystic disease in mice^32,33^. However, the uniqueness of our approach is that, rather than transactivation, it relies on preventing inhibition, making it unlikely that *PKD1* will rise to the supratherapeutic range.

Our studies clarify the role of *PKD1* in cyst initiation and its continual expansion. A unifying and parsimonious explanation for ADPKD onset is that cystogenesis ensues when the *PKD1* dosage falls by 70-80%, dipping below a critical threshold^3^. Thus, heterozygous germline *PKD1* inactivation alone cannot account for this magnitude of dose reduction. Additional stochastic events that repress the remaining allele are required and critical in determining disease onset. However, the somatic inhibitory mechanism(s) in addition to the ‘second-hit’ mutations are unknown. Our discovery of cis-interference resulting in inefficient translation of mRNAs transcribed by the non-inactivated *PKD1* allele represents a novel and targetable ADPKD onset mechanism. Along with cyst initiation, *PKD1* inhibition unleashes large-scale transcriptomic and metabolic dysregulation and activates numerous oncogenic pathways, such as cAMP and c-Myc/Yap^4,34–42^. In turn, this downstream cyst-pathogenic signaling fuels cyst expansion. Remarkably, even in the face of such widespread dysregulation, a recent elegant study reported that transgenic *Pkd1* or *Pkd2* reconstitution rapidly reverts established cystic disease in mice^43^. Consistently, we found that acute *Pkd1/2* derepression reigns in established cystic disease and makes *Pkd1*-mutant cells resistant to pro-cystogenic stimuli such as cAMP and SAM. Collectively, these observations point to *PKD1* as the primary, if not the sole, factor governing both cyst onset and growth.

We report an unexpected finding that *Pkd2* influences the cystic phenotype of *Pkd1*-mutant models. PC1 and PC2 physically interact and are coexpressed at multiple subcellular locations, indicating that the two proteins function in the same physiological pathway^44–47^. We add a new dimension by extending this relationship into the pathological context. Perhaps, enhancing *Pkd2* expression in *Pkd1*-mutant cells may lead to improved PC1 trafficking and more heteromeric PC1-PC2 protein complexes.

Our work highlights an underappreciated aspect of gene regulation. Canonical thinking is that miRNAs simultaneously but subtly repress large mRNA networks. In contrast, we made the surprising finding that, under certain circumstances, miRNAs transform into potent single-target inhibitors. In fact, we show that an individual 3’-UTR MBE has a marked disease-modifying impact. Most miRNAs are dispensable for homeostatic tissue functions, and they are pharmaceutically inhibited with relative ease. Despite these favorable characteristics, miRNA-based drug development has languished in comparison to other forms of RNA therapeutics^48^. This is partly because the pleiotropic molecular mechanism of numerous downstream mRNA targets makes it difficult to validate the miRNA biological effect or develop pharmacodynamic readouts of anti-miRNA drugs. We argue that prioritizing miRNAs that function as tonic inhibitors of a handful of disease-central mRNAs is likely to be a fruitful drug development strategy. Importantly, our insights are transferable, and we speculate that similar modes of therapeutically targetable cis-inhibitory regulation exist in other disorders, especially haploinsufficient monogenetic conditions.

## MATERIALS AND METHODS

### Generation of 3’-UTR cell lines via CRISPR/Cas9

We deleted the miR-17 binding site from the *Pkd1* or *Pkd2* 3’-UTR using CRISPR/Cas9. We designed sgRNAs using https://www.benchling.com and ordered the sequences from IDT. The sgRNA pair targeted DNA sequences upstream and downstream of the miR-17 motif in the *Pkd1* or *Pkd2* genes. We cloned the sgRNAs into the CRISPR mammalian expression vector pSpCas9(BB)-2A-GFP^49^. Using these sgRNA encoding plasmids, we generated the *Pkd1*^*RC*Δ17/-^ cell line and the *Pkd1*^*RC*/-^; *Pkd2*^Δ17/Δ17^ cell line as described next. To generate the *Pkd1*^RCΔ17/-^ cell line, we transfected *Pkd1*^RC/-^ cells with 0.6 μg of the SpCas9-2A-GFP plasmid carrying the upstream or the downstream sgRNA using Lipofectamine 3000. After 72 hours, we performed FACS to select GFP-positive cells with the top 5% intensity. These cells underwent clonal expansion in 96-well plates. Well-formed colonies were screened for the absence of the miR-17 binding site by DNA PCR of the targeted *Pkd1* genomic sequence. Clones with expected deletion bands were confirmed by Sanger sequencing. Two *Pkd1*^*RC*Δ17/-^ clonal cell lines with confirmed deletions were further characterized and analyzed along with their parental control cell lines, as shown in **Fig. 2** and **Suppl. Fig. 4**. We used the same strategy and experimental approach for generating the two *Pkd1*^*RC*/-^; *Pkd2*^Δ17/Δ17^ cell lines (**Fig. 4** and **Suppl. Fig. 8**). The sgRNA sequences and genotyping primers are provided in **Supplementary Table 3**.

### Generation of 3’-UTR mice via CRISPR/Cas9

The following strains of mice were used: (1) for the mouse models shown in **Fig. 1**, wild-type C57BL/6N female and male mice were used; (2) for the mouse models shown in **Figs. 2** and **4**, we used *Ksp*^Cre^; *Pkd1*^RC/RC^ mice maintained on a C57BL/6J background by our laboratory. Prepubertal female mice underwent superovulation using a standard hormone regimen. The epididymis was collected from male mice for sperm harvest. After in vitro fertilization, one-cell fertilized eggs were isolated. CRISPR reagents (IDT) were delivered to the cytoplasm via electroporation using a Nepa21 Super Electroporator (NEPAGENE, Ichikawa, Japan). The eggs that survived the electroporation were washed and cultured in fresh M16 media in microdrop cultures. The eggs were then surgically transferred into the oviducts of day 1 pseudopregnant ICR females. At 21 days of age, founder mice were screened for deletion of the miR-17 binding site by genotyping, and confirmation of deletion was performed by Sanger sequencing.

### ADPKD mouse models

*Ksp*^Cre^, *Pkd1*^F/F^, and *Pkd1*^RC/RC^ mice have been previously described^4,23,28,50^. All mice were maintained on a C57BL/6J background. At prespecified time points, mice were anesthetized using an approved protocol, and blood was obtained via cardiac puncture. The right kidney was weighed to obtain the KW/BW ratio and immediately flash frozen for future molecular analysis. The left kidney was perfused with ice-cold 1X PBS and 4% (wt/vol) paraformaldehyde. The kidney was subsequently paraffin-embedded. All studies used equal numbers of males and females. The UT Southwestern Institutional Animal Care and Use Committee approved all experiments involving animals.

### Histology

Tissue embedding in paraffin and subsequent sectioning were performed using standard protocols by the Histology core at UT Southwestern Medical Center. The tissues were cut into 5 μm sections and stained with hematoxylin-eosin (H&E) for histological analysis. The stained sections were imaged using a slide scanner.

### RNA

A Qiagen miRNEASY kit was used for total RNA extraction. cDNA was prepared using an Invitrogen First Strand Superscript III cDNA synthesis kit. Q-PCR was performed using iQ SYBR Green Supermix (Bio-Rad). All samples were loaded in duplicate or triplicate on the CFX ConnectTM Real-time PCR detection system. 18s was used to normalize mRNA expression. The sequences of the primers are shown in **Supplementary Table 4**.

### PC1 and other Western blots

Total protein was isolated from kidneys or cells using lysis buffer made from mixing T-PER tissue protein extraction reagent (Invitrogen, catalog# 78510) with a protease phosphatase inhibitor tablet (Fisher, catalog# PIA32961) according to the manufacturer’s instructions. The lysis buffer was prepared and stored as one-time aliquots at −80°C. The aliquots were thawed on ice immediately before protein isolation. Protein concentration was measured using the Bradford Assay reagent. Protein samples were prepared in 4X NuPAGE LDS Sample Buffer with 0.5% b-mercaptoethanol (Sigma, catalog# M6250) for all proteins except for PC1 and PC2 and their loading control beta-actin, which were prepared with 0.1 M DTT (Sigma, catalog# D0632). The samples were always freshly prepared before gel electrophoresis. BME samples were boiled for 5 minutes at 98°C before loading on gels. The DTT samples were incubated at 25°C for 10 minutes before loading on gels.

For PC1 detection, the samples were run on the NuPAGE™ 3-8% Tris-Acetate Protein Gel (Invitrogen, EA03785) at 160 V for 1.5 hours on ice. A high molecular weight protein ladder (Invitrogen, catalog# LC5699) was used in each gel to track 460 kDa proteins. Electrophoretically separated proteins were transferred using the Invitrogen transfer system at 200 mAmps for 100 minutes on ice or at 4°C. To detect other proteins, the samples containing 10 μg of protein were run on mini-PROTEAN SDS-polyacrylamide precast gels. A standard molecular weight ladder was used in each gel to track protein sizes. The gels were run at 150 V until the dye ran out. The proteins were transferred to a nitrocellulose membrane using the Trans-Blot Semi-Dry Transfer system on the mixed MW program.

After completing the transfer, the membranes were blocked with 5% fat-free milk and probed overnight at 4°C with primary antibodies. The membranes were washed three times with 1x TBS-Tween the next morning before and after probing for one hour with a secondary antibody. Goat-anti-rabbit or anti-mouse HRP-conjugated IgG was used as the secondary antibody. HRP-conjugated actin antibody (Sigma, catalog# a3854) was used to measure total protein. The blots were developed using the chemiluminescence substrate SuperSignal West Dura, ECL, or Femto from Pierce. The blots were developed using the Bio-Rad digital imager. The protein bands were quantified using Imagelab software from Bio-Rad. Each Western blot was repeated at least three times. Ten micrograms of protein from cells or kidneys was run on gels for the detection of < 150 kDa proteins. A total of 40-60 μg of protein was run on gels to detect heavy molecular weight (462 kDa) full-length PC1 protein. All the primary antibodies were used at a 1:1000 dilution, except for PC1 (used at 1:500), and the secondary antibodies were used at a 1:5000 dilution. The following primary antibodies were used: PC1 (7E12 Santa Cruz, catalog# sc-130554); PC2 (gift from the Baltimore PKD Core); pCREB (Cell Signaling, catalog# 9198); c-Myc (Abcam, catalog# ab185656), YAP1 (Cell Signaling, catalog# 4912); Mettl3 (Invitrogen, catalog# MA5-27527).

### Immunofluorescence on tissue samples

Paraffin sections of kidney tissues were used for immunofluorescence staining. Briefly, the slides were deparaffinized by first baking at 60°C for 1 hour and then washing in Histo-clear (Fisher, HS-2001) three times for 5 minutes each. Next, the slides were re-hydrated through 100%, 95%, and 70% ethanol washes before incubation in 1X PBS. The slides were then subjected to antigen retrieval with sodium citrate. The slides were treated with sodium borohydride to quench autofluorescence for 40 min. The slides were washed in 1X PBS three times and then blocked in 1X PBS+10% goat serum+0.1% BSA (antibody block) for at least 1-2 hours at RT. Sections were incubated with primary antibodies overnight. Primary antibodies were diluted with antibody block at a 1:500 dilution. Slides were washed in 1X PBS three times for 5 minutes each, treated with Alexa Fluor secondary antibodies (diluted using the antibody block to a 1:500 dilution) for 1 hour, and then washed three times for 5 minutes each. The slides were mounted using Vecta Shield containing Dapi. The slides were imaged using the Zeiss Compound Light microscope or the Zeiss Axioscan Z1 slide scanner. The following antibodies were used: DBA (Vector Labs, catalog# B-1035), THP (Biomedical Technologies, catalog# BT-590), LTA (Vector Labs, catalog# B-1325), MRC1 (Abcam, catalog# ab64693), pCREB1 (Cell Signaling, catalog# 9198), and pHH3 (Sigma, catalog# H0412). Processing, immunostaining, and imaging of slides were performed simultaneously within each experiment.

### Immunofluorescence on cells

Immunofluorescence staining was performed on cells grown on 8-chambered slides (Fisher, catalog# 154534PK). The cells were fixed with 100% ice-cold methanol for 5 minutes at 4°C. The slides were washed with 1X PBS 3 times for 5 minutes. The cells were then blocked in 1X PBS+10% goat serum+0.1% BSA+0.1 M glycine+0.1% Tween 20 (antibody block) for at least

30 minutes at room temperature. Primary antibodies were diluted with antibody block at a 1:100 dilution and added to the slides for 2 hours. Slides were washed in 1X PBS three times for 5 minutes each, treated with Alexa Fluor secondary antibodies (diluted using the antibody block to a 1:500 dilution) for 1 hour, and then washed three times for 5 minutes each. The slides were counterstained in DAPI (Fisher, catalog# ICN15757410) diluted at 1:10000 in distilled water for 10 minutes before imaging under a Zeiss Compound Light microscope. For each experiment, the control and treatment cells or the control and the Δ17 cells were seeded simultaneously on different chambers of the same slide. Processing, immunostaining, and imaging of slides were also performed simultaneously.

### MitoTracker analysis

MitoTracker Red CMXRos (Thermo Fisher, catalog# M7512) was used to analyze the mitochondrial membrane potential in live cells. The lyophilized MitoTracker® product was dissolved in dimethylsulfoxide (DMSO) to a final concentration of 1 mM and stored at −20°C in small aliquots. Cells grown to 40-70% confluency were washed with sterile PBS and then treated with regular DMEM serum-free media containing 100 nM MitoTracker for 8 minutes. Immediately thereafter, the media was replaced with regular growth media and imaged under a Zeiss Compound Light microscope. The images were taken at the same exposure time for the samples of the same experiment. The intensity of fluorescence is directly proportional to membrane potential.

### 3D Cystogenesis Assay

25 μl of 100% Matrigel (Fisher, catalog# 354234) was spread onto each well of an 8-chambered slide with precooled 200 μl sterile pipette tips. The plate was then placed in a 37°C incubator for 30 minutes for the Matrigel to set. In the interim, cells were trypsinized, washed once with PBS, filtered through a 40 μm cell strainer to create a single-cell suspension, and counted. Cells were seeded on the Matrigel-coated slide at a seeding density of 5000 cells/well in a 300 μl volume of growth media containing 2% Matrigel. For each cell line or treatment condition, cells were seeded in triplicate and incubated at 37°C for 7 days to allow for the growth of 3D cysts in suspension. During this time, the wells were supplemented with growth medium 72 hours after initial placement into Matrigel. On day 7, the chamber slides were imaged on a Leica DMI 3000B light microscope. The images were analyzed using ImageJ software to obtain cyst size measurements. Each assay was repeated at least three times. Measurements from each experiment were combined and analyzed for statistical significance.

### *Ex vivo* Organ Culture

Female mice bearing potential *Pkd1*^+/+^; *Pkd1*^Δ/+^ and *Pkd1*^Δ/Δ^ embryos were dissected at embryonic day (E) 13.5 in PBS to harvest the kidneys and tail. The tail of each embryo was used for DNA extraction and subsequent genotyping. The kidneys were set up for culture on Whatman membranes (Sigma, catalog# WHA110409) in an air-medium interface as described^26^. The kidneys were cultured in basal DMEM (Thermo Fisher, catalog# 12500) containing 10% fetal bovine serum (FBS), 2% PenStrep (Invitrogen, catalog# 1514022), 5 μg/ml insulin, 5 μg/ml transferrin, 2.8 nM sodium selenite, 25 ng/ml prostaglandin E and 32 pg/ml T3. One kidney was grown in the above media, and the contralateral kidney was grown in 100 μM 8-Br-cAMP (Sigma, catalog# B7880)-supplemented media. Using a second cohort of mice, one kidney was grown in 100 μM 8-Br-c-AMP or 100 μM 8-Br-c-AMP + 250 μM SAM. For all the cultures, the media was changed every 48 hours. The cultures were imaged live using the Zeiss Stereo Lumar microscope on day 4. The cysts were measured and analyzed using ImageJ software. At the end of 6 days, the kidneys were flash-frozen and stored at −80°C until further use for RNA or protein extraction.

### alamarBlue Assay

*Pkd1*^RC/-^ and *Pkd1*^RCΔ17/-^ cells (3 × 10^3 density) were seeded on 96-well plates. The next morning, the medium was changed to contain 1X alamarBlue reagent (Invitrogen, catalog# DAL1025) and vehicle, 100 μM 8-Br-cAMP, 100 μM SAM, or 17 mM glucose. Colorimetric readings were taken at 570 nm and 600 nm in a microplate reader after 12 hours. The redox reaction of alamarBlue was used to assess cell proliferation quantitatively. N=8 was used for each condition. The values were plotted as a scaled heatmap using the Python MatplotLib package. The same experimental approach was used for the *Pkd1*^RCΔ/-^; *Pkd2*^Δ17/Δ17^ cells and the control cells *Pkd1*^RC/--^; *Pkd2*^+/+^.

### Serum Electrolytes

Serum creatinine was measured by capillary electrophoresis by the UT Southwestern O’Brien Center. BUN was measured by Vitros250 Analyzer by the UT Southwestern Metabolic Phenotyping Core.

### RNA-seq preprocessing

Sequencing quality control was performed with FastQC v0.11.8. RNA-seq reads were trimmed, and low-quality reads were removed using Trimgalore v0.6.3_dev (https://www.bioinformatics.babraham.ac.uk/projects/trim_galore/) with the “paired” parameter and length of 150 bps. Trimmed fastq sequences were aligned to the mouse reference genome GRCm38 using STAR aligner v2.5.3a with the produced bam files sorted by coordinate by using the option “-- outSAMtype BAM SortedByCoordinate. “^51,52^ Raw read gene counts were obtained using STAR aligner with the options “—quantMode GeneCounts” and “--sjdbGTFfile” with gene models in GTF format obtained from mouse EnsEMBL release 94. Alignment quality control and read mapping statistics were obtained from Picard tools v2.20.3 using the function “CollectMultipleMetrics” (http://broadinstitute.github.io/picard/).

### RNA-seq data analysis

Raw gene counts were used for quality control and differential expression analysis. Raw counts were normalized to the total number of reads by calculating log2CPM (counts per million), and lowly expressed genes (average Log2CPM < −3) were eliminated before differential gene expression analysis. TPM (transcript per million) quantification was performed using RSEM v1.3.1, and differential gene expression analysis was performed using the limma-trend (version 3.40.6) in R^53,54^.

### Transcript quantification

Individual *Pkd1* transcript quantification was performed using Salmon v1.3.0^55^. The five different transcript versions for *Pkd1* were added to the RefSeq mm9 fasta reference transcriptome to build a novel Salmon index. Then, the fastq files were directly mapped and read counts, and TPM values were quantified with the standard process.

### *In vitro* RGLS4326 experiments

*Pkd1*^RC/-^ cells were seeded at 2 × 10 ^5 confluence in 6-well plates. The next morning, cells were transfected using Lipofectamine 3000 with a vehicle, control oligonucleotide, or RGLS4326 at a final concentration of 100 μM. Forty-eight hours after transfection, the cells were collected for RNA extraction. Seventy-two hours after transfection, cells were harvested for protein or further seeded for alamarBlue assay and 3D cystogenesis assay. For the experiment shown in Figure 5C, the 3D cystogenesis assay was performed with untreated *Pkd1*^RC/-^ cells, as described in the methods section of the 3D cystogenesis assay with the following changes. On day 4 of Matrigel culture, the wells were imaged using the Leica light microscope DMI 3000B, and then, the cultures were transfected with a vehicle, 100 μM control oligonucleotide, or 100 μM RGLS4326 and grown for 3 additional days. On day 7, the samples underwent imaging to assess cyst size.

### RGLS4326 mouse experiments

The Ksp^Cre^; *Pkd1*^F/RC^ mouse line was used for the drug studies. Mice were randomly assigned and administered 20 mg kg^−1^ vehicle (PBS), control oligonucleotide, or RGLS4326 via subcutaneous injections. For the first cyst prevention study (**Figure 5D-G**), mice were injected on postnatal days (P) 10, P11, P12, and P16 and sacrificed on P18. Nontransgenic strain-matched mice were also sacrificed on the same days. For the second disease stabilization study (**Figure 5H-K**), mice were injected at P16 and P17 and sacrificed at P26. One mouse from the study succumbed to the disease and died earlier than 26 days of age. For the third long-term study (**Figure 5L-N**), mice were injected on P16 and P17 and then every week until 18 weeks of age. Another cohort of mice received the same dose of RGLS4326 treatment on P16 and P17 and then semimonthly thereafter until 18 weeks of age. The mice were observed every day for 18 weeks to note death. At the end of 18 weeks, the surviving mice were sacrificed to harvest tissue. Equal numbers of males and females were used in all study groups.

### Human ADPKD cell experiments

Primary human ADPKD cyst cells were obtained from PKD Research Biomarker and Biomaterial Core at the University of Kansas Medical Center (KUMC). The use of surgically discarded kidney tissues complied with federal regulations and was approved by the Institutional Review Board at the University of Kansas Medical Center. Each primary cell line was cultured in DMEM/F12++ (Gibco, catalog# 10565–018) supplemented with 10% FBS, 5 μg kg^−1^ insulin, 5 μg mL^−1^ transferrin, and 5 ng mL^−1^ sodium selenite and incubated in an atmosphere of 95% air and 5% CO2 at 37 °C until 80% confluency. At the 2^nd^ passage, cells from each human donor underwent reverse transfection using CRISPRMAX reagent (Invitrogen) containing Cas9 protein (IDT) and synthetic sgRNAs (IDT) or were transfected with vehicle (lacking Cas9). Cas9- or vehicle-transfected cells (control) were then seeded into 6-well plates and chamber slides. After 72 hours, the cells were harvested for genotyping, Western blot analysis, and immunofluorescence/MitoTracker staining. In addition, at 72 hours posttransfection, cells were trypsinized and plated at 4000 cells/well density in 130 μl of media plus Matrigel (Corning, catalog# 354234) in a 96-well plate (Corning, catalog# 353072). Media was replenished 72 hours after the initial placement into Matrigel. Cyst images were obtained on the 7^th^ day of Matrigel culture (10^th^ day after Cas9/SgRNA or vehicle transfection). One hundred cyst images were obtained for Cas9- or vehicle-transfected cells from each donor. Similarly, 72 hours posttransfection, cells were seeded at 2000 cells/well density in 96-well plates for the alamarBlue proliferation assay. The following day, the media was replaced with growth media containing 1X alamarBlue, and readings were taken 12 hours later.

### Statistical Analysis

Student’s t-test was used for pairwise comparisons or analysis of variance (ANOVA), followed by Tukey’s post hoc test for multiple comparisons. The Mantel-Cox test was used for the analysis of mouse survival. *P* <0.05 was considered statistically significant. The sample size and *P* values are mentioned in the figure graphs, the figure legends, and the results section. For *in vivo* experiments, *N* is the number of mice analyzed. For *in vitro* experiments, *N* refers to the number of biological replicates. For the RGLS4326 studies, animals were randomly assigned to treatment arms. Investigators were not blinded to the treatment or the genotypes of the animals.

## Data Accessibility

The RNA-Seq data are accessible.

## Acknowledgments

The work was supported by grants from the National Institutes of Health (R01DK102572) and the Department of Defense (D01 W81XWH1810673) VP. RL is supported by the National Institute of Health (K08DK117049) and grants from the PKD Foundation and the American Society of Nephrology KidneyCure Grants Program. We thank the O‘Brien Kidney Research Core Center (P30DK079328) at UT Southwestern Medical Center (UTSW), the Eugene McDermott Center for Human Growth and Development Sequencing Core and Bioinformatics Lab at UTSW, the UTSW Molecular Pathology Core, and the Whole Brain Microscopy Facility at UTSW, Genewiz, and Monoceros for providing critical reagents and services. Human ADPKD and normal human kidney cells and tissues were provided by the PKD Biomarkers and Biomaterials Models Core, located at the University of Kansas Medical Center. The Core is part of the PKD Research Resource Consortium, supported by the National Institutes of Health/NIDDK (U54 DK126126).

## SUPPLEMENTARY INFORMATION

**Supplementary Figure 1:**
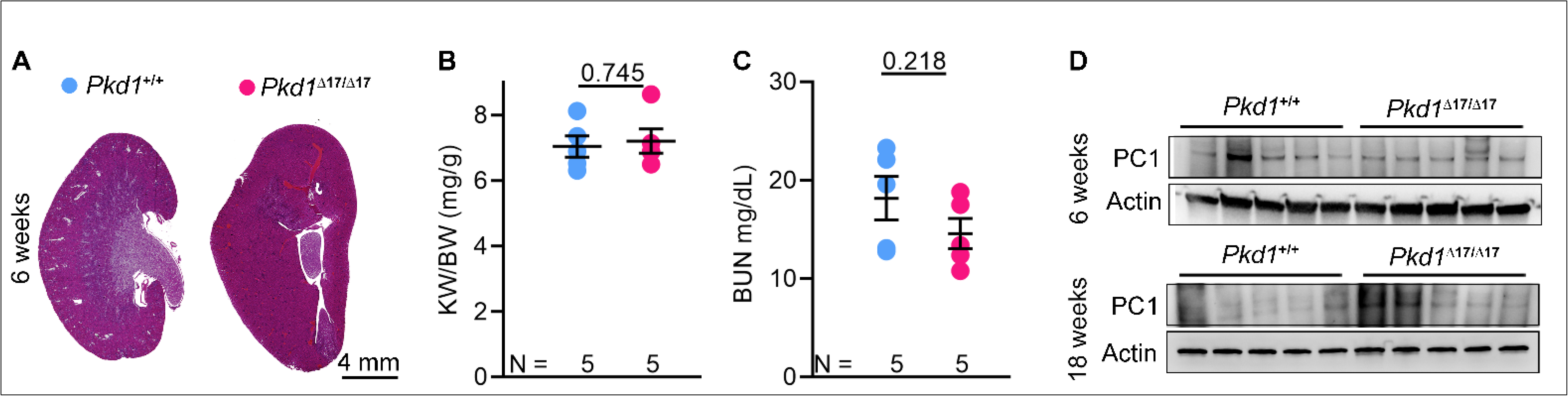
Characterization of *Pkd1*^Δ17/Δ17^ mice. **A.** H&E-stained kidney sections of 6-week-old *Pkd1*^+/+^ and *Pkd1*^Δ17/Δ17^ mice are shown. Both mice exhibited normal kidney histology. **B & C/** Kidney-weight-to-body-weight (KW/BW) ratio and BUN levels of 6-week-old *Pkd1*^+/+^ and *Pkd1*^Δ17/Δ17^ mice are shown. **D.** Immunoblot showing equivalent PC1 expression in the kidneys of 6- or 18-week-old *Pkd1*^+/+^ and *Pkd1*^Δ17/Δ17^ mice. Error bars indicate SEM. Student’s t-test was used for statistical analysis (B-C).

**Supplementary Figure 2:**
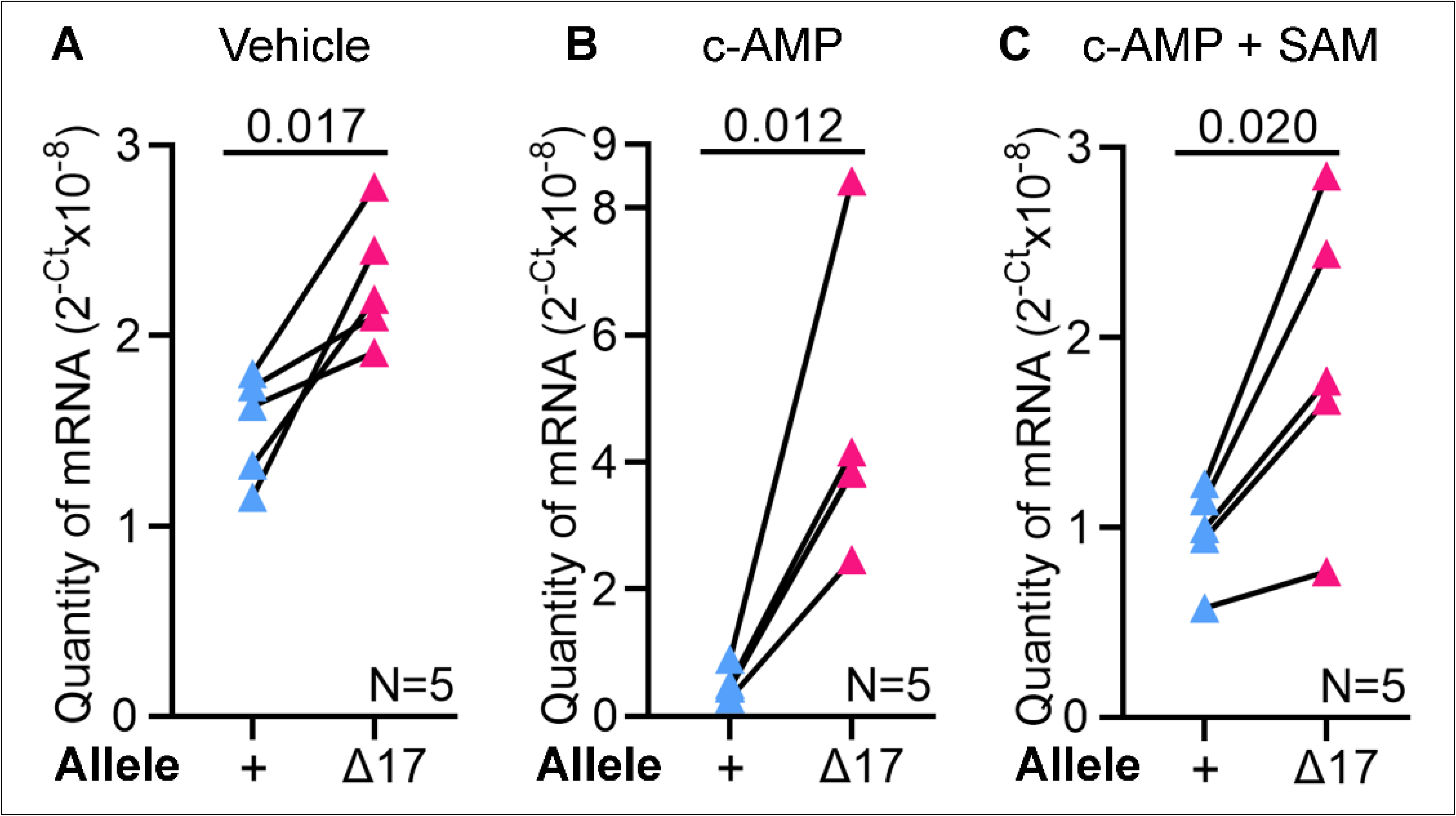
*Pkd1* is cis-inhibited via its miR-17 3’-UTR motif. Allele-specific qRT-PCR analysis showing the quantity of *Pkd1* mRNAs produced by the wild-type (+) and Δ17 alleles in *ex vivo* kidney cultures of *Pkd1*^Δ17/+^; mice treated with vehicle (**A**), c-AMP (**B**), or c-AMP plus SAM (**C**). The *Pkd1*^Δ17^ allele produced more mRNA transcripts than the *Pkd1*^+^ allele. This difference was even more pronounced in the presence of c-AMP. Statistical analysis: paired t-test. Error bars indicate SEM.

**Supplementary Figure 3:**
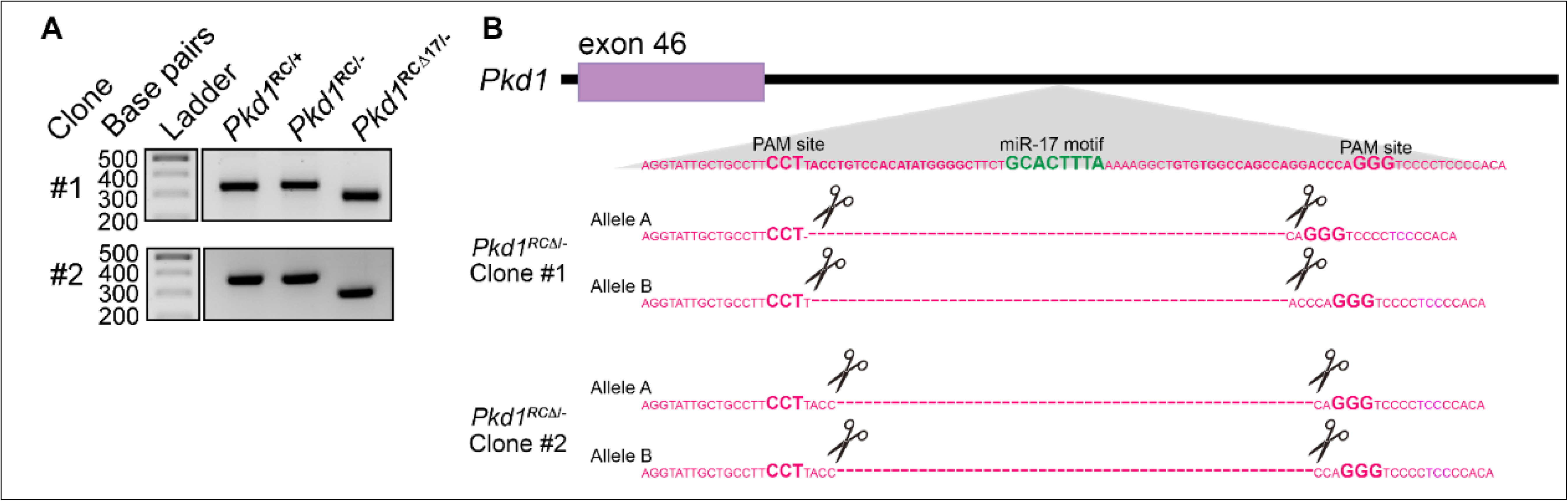
Characterization of CRISPR-edited *Pkd1*^*RC/-*^ cell lines. **A.** PCR products obtained after amplifying the DNA (encoding the *Pkd1* 3’UTR segment) from parental and CRISPR-edited cell lines. The lower band indicates the Δ17 genotype. **B.** Graphical illustration of Sanger sequencing results from the Δ17 bands of each clone confirm deletion of the miR-17 motif from both *Pkd1* alleles.

**Supplementary Figure 4.**
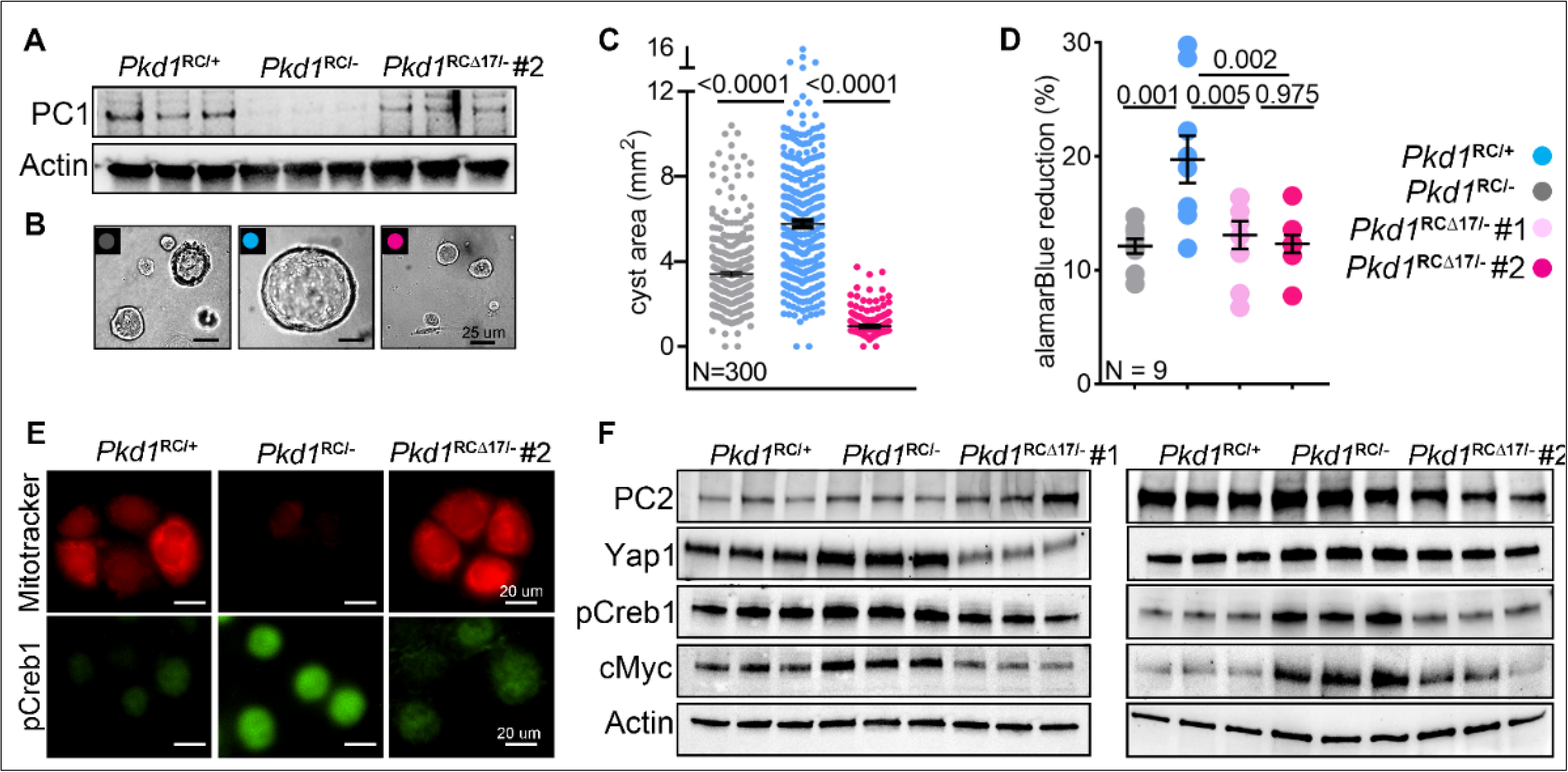
Phenotypic characterization of CRISPR-edited *Pkd1*^*RCΔ17*^ cell lines. **A.** Western blot showing PC1 derepression in *Pkd1*^RCΔ17/-^ clone #2 compared to the *Pkd1*^RC/-^ parent cell line. Actin was used as a loading control. **B, C.** 3D cyst images and quantification showing a substantial reduction in cyst size of *Pkd1*^RCΔ17/-^ cells compared with *Pkd1*^RC/-^ cells. **D.** AlamarBlue assay showing reduced proliferation of *Pkd1* clones that lack the miR-17 motif at 12 hours compared to the parental cell line. **E.** MitoTracker images and IF staining for pCreb1 showing restored mitochondrial membrane potential (red) and reduced pCreb1 (green) expression in *Pkd1*^RCΔ17/-^ clone #2 compared to the parental cell line. **F.** Western blot characterization of both *Pkd1*^RCΔ17/-^ cell lines showing reduced expression of the cyst-promoting genes Yap1, pCreb1, and c-Myc compared to parental *Pkd1*^RC/-^ cells. As a pertinent control, PC2 expression remained unchanged. Actin served as the loading control. Error bars indicate SEM. Statistical analysis: one-way ANOVA, Tukey’s multiple comparisons test (C, D).

**Supplementary Figure 5.**
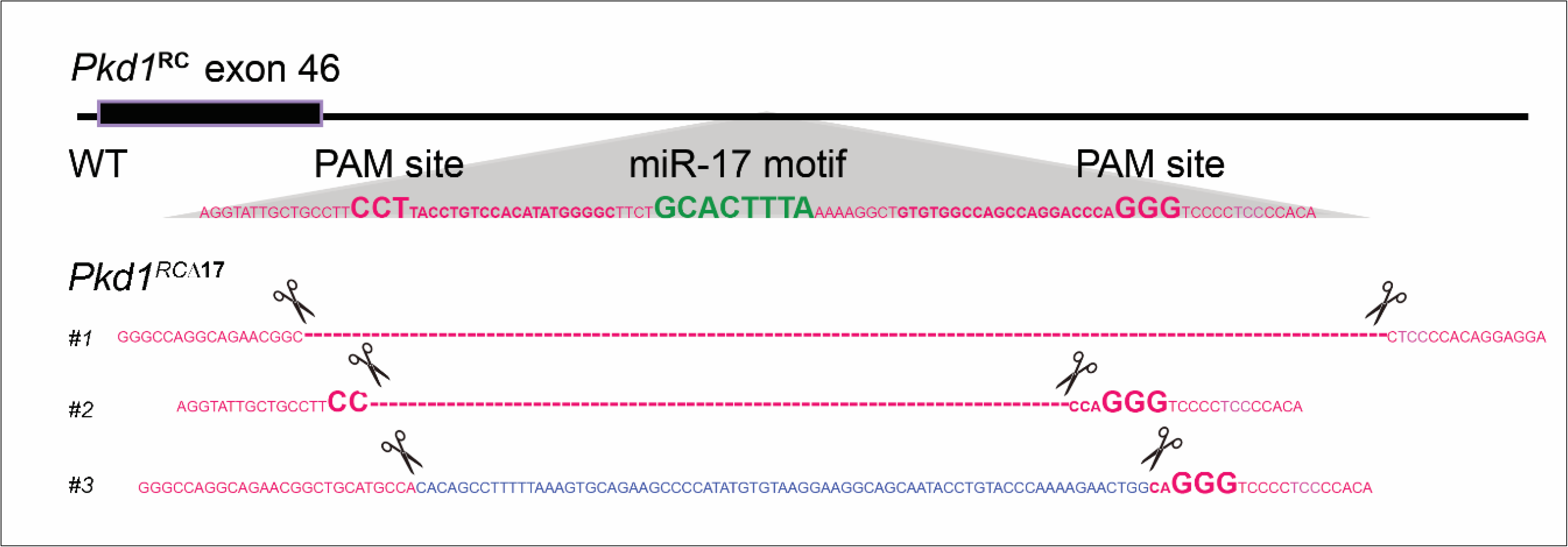
Characterization of CRISPR-edited *Pkd1*^RC/RC^ mice. Graphical illustration of Sanger sequencing from tail DNA from the three CRISPR-edited founders. Founders #1 and #2 harbor 108 bp and 53 base pair deletions, respectively, including the miR-17 motif. Founder #3 also lacked the miR-17 motif but acquired a 72 base pair insertion (blue), resulting in a net loss of 18 base pairs in the 3’-UTR sequence.

**Supplementary Figure 6:**
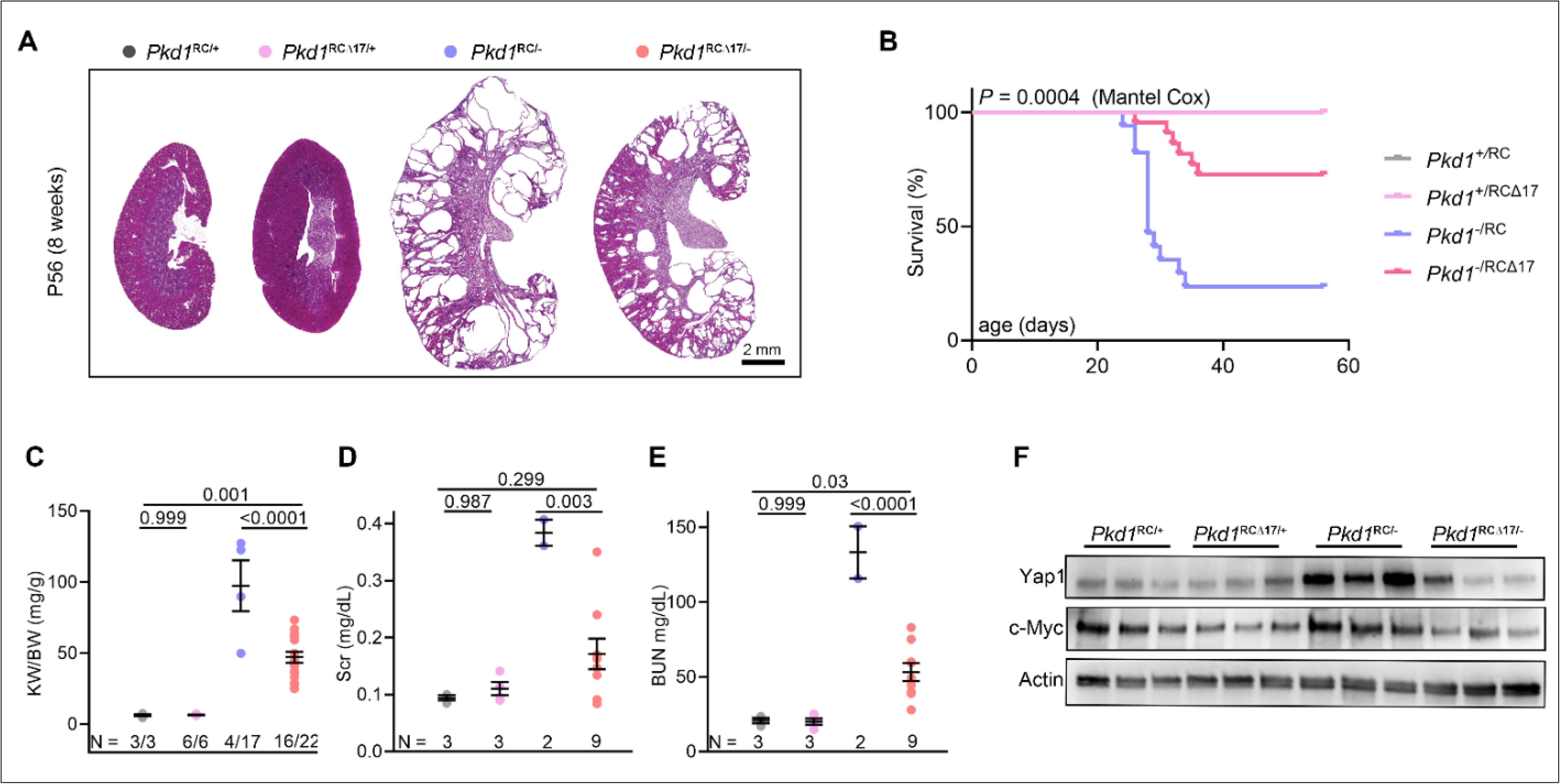
Monoallelic *Pkd1* derepression suppresses disease progression. A cohort of progeny derived from founder #2 was prospectively monitored until 8 weeks of age. **A.** H&E-stained kidney sections of mice of the indicated genotypes that survived until 8 weeks of age are shown. **B.** Kaplan-Meir survival curves of mice with the indicated genotypes are shown. **C-E.** KW/BW, serum creatinine (Scr), and BUN levels of the surviving 8-week-old mice with the indicated genotypes are shown. **F.** Immunoblots showing Yap1 and c-Myc expression in the kidneys of 18-day-old mice with the indicated genotypes. Actin served as the loading control. Error bars indicate SEM. Statistical analysis: one-way ANOVA, Tukey’s multiple comparisons test (C-E); log-rank Mantel-Cox (B)

**Supplementary Fig 7:**
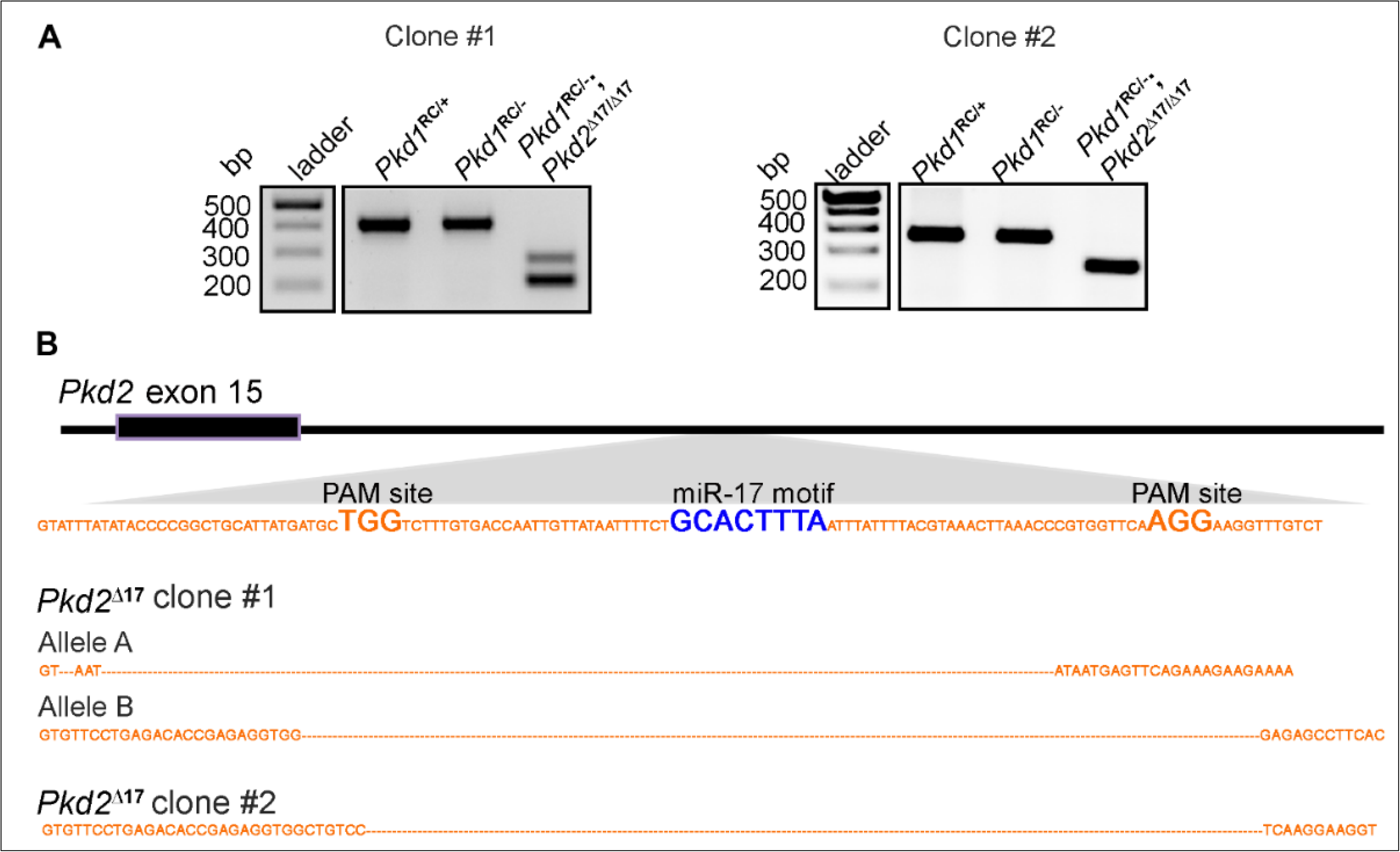
Characterization of *Pkd1*^*RC/-*^ cell lines lacking the miR-17 motif in the *Pkd2* 3’-UTR. **A.** PCR products obtained after amplifying the DNA (encoding the *Pkd2* 3’UTR segment) from parental and CRISPR-edited cell lines. The lower bands indicate the Δ17 genotype. **B.** Graphical illustration of Sanger sequencing results from the Δ17 bands of each clone confirm deletion of the miR-17 motif from both *Pkd2* alleles.

**Supplemental Figure 8.**
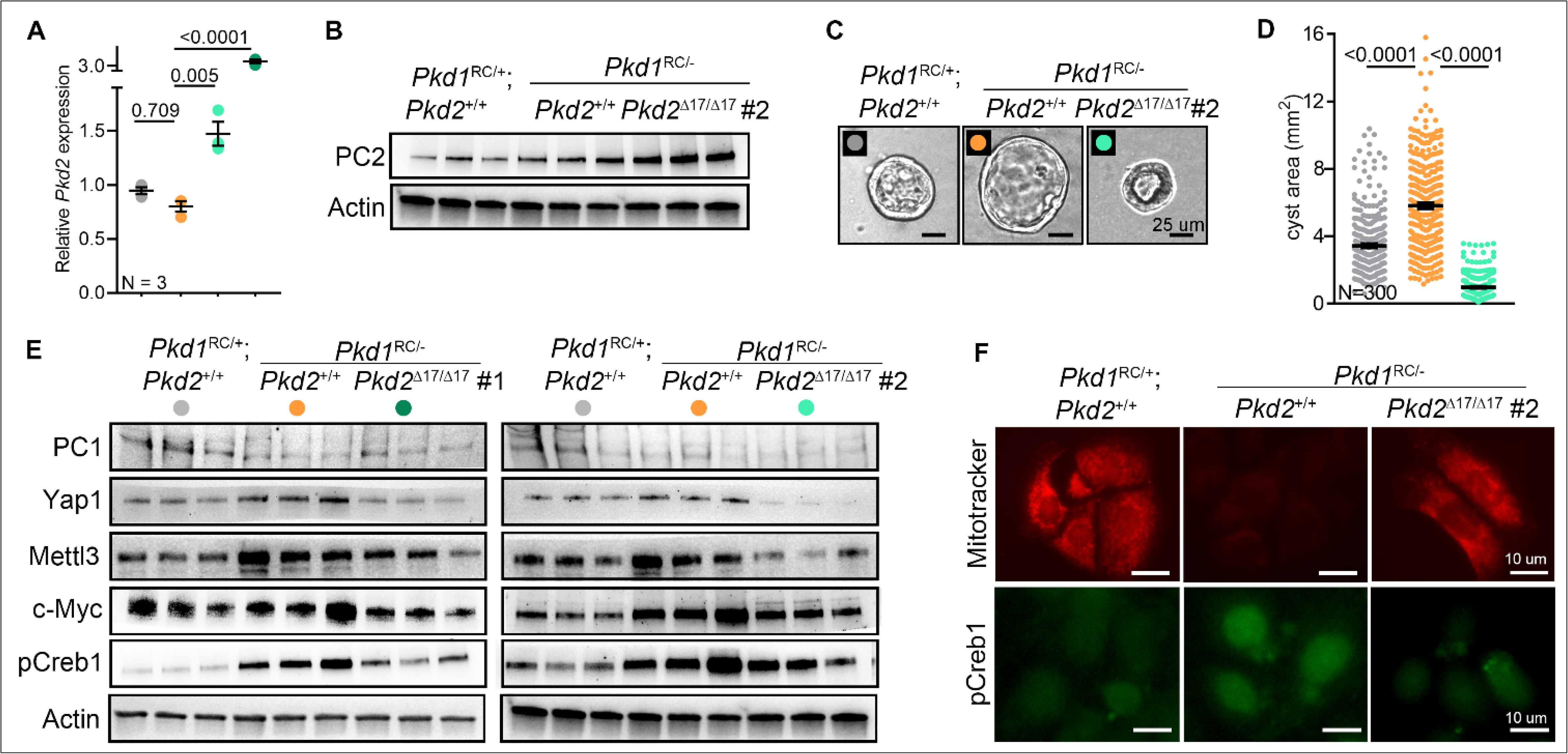
Phenotypic characterization of CRISPR-edited *Pkd1*^*RC/-*^; *Pkd2*^Δ17/Δ17^ cell lines. **A, B.** qRT-PCR and Western blot showing *Pkd2* and Polycystin-2 derepression, respectively, in both *Pkd1*^RC/-^; *Pkd2*^Δ17/Δ17^ clones #1 and #2 compared to the parental *Pkd1*^RC/-^ cell line. **C-D** Representative images and quantification of 3D cyst assay comparing indicated cell lines showing marked reduction in cyst size of *Pkd1*^*RC/-*^; *Pkd2*^Δ17/Δ17^ clone #2 compared to parent *Pkd1*^RC/-^ cell line. **E.** Western blot characterization showing reduced expression of the cyst-promoting genes Yap1, Mettl3, c-Myc, and pCreb1 in both *Pkd1*^RC/-^; *Pkd2*^Δ17/Δ17^ clones compared to the parent *Pkd1*^RC/-^ cell line. Pertinently, PC1 expression remained unchanged. Actin served as a loading control. **F.** MitoTracker images and IF staining for pCreb1 showing restored mitochondrial membrane potential (red) and reduced #2 compared to the parental cell line. Error bars indicate SEM. Statistical analysis: one-way ANOVA, Tukey’s multiple comparisons test (A, D).

**Supplementary Figure 9.**
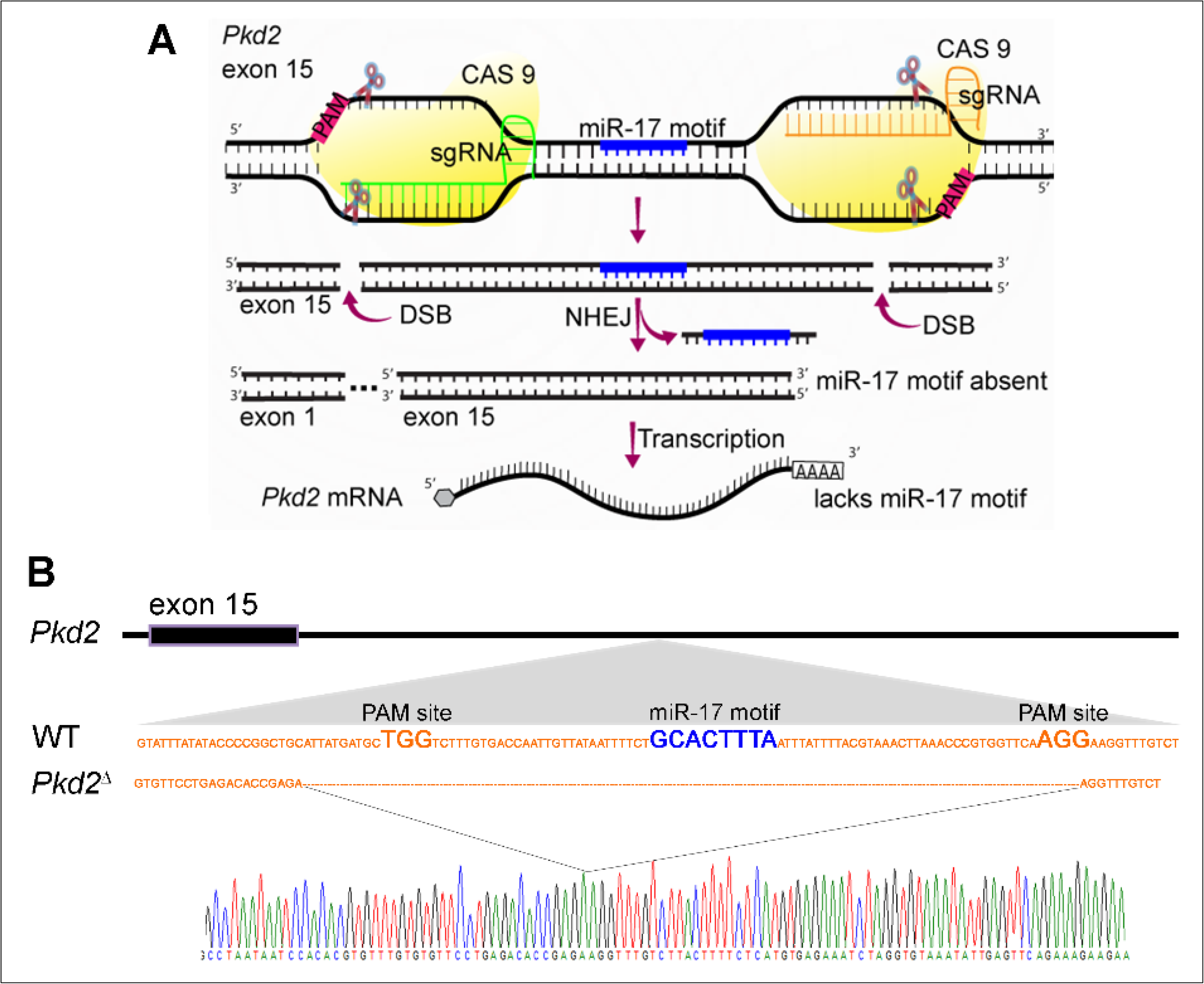
CRISPR editing and characterization of *Pkd2*^Δ17^ mice that lack the miR-17 motif in the *Pkd2* 3’-UTR. **A.** Graphic illustrates the CRISPR-editing strategy used to remove the miR-17 motif from exon 15 of *Pkd2*. **B.** Graphical illustration and Sanger sequencing results of PCR products from the tail DNA of founder mice showing 139 base pair deletions from the *Pkd2* 3’-UTR, including the miR-17 motif.

**Supplementary Figure 10:**
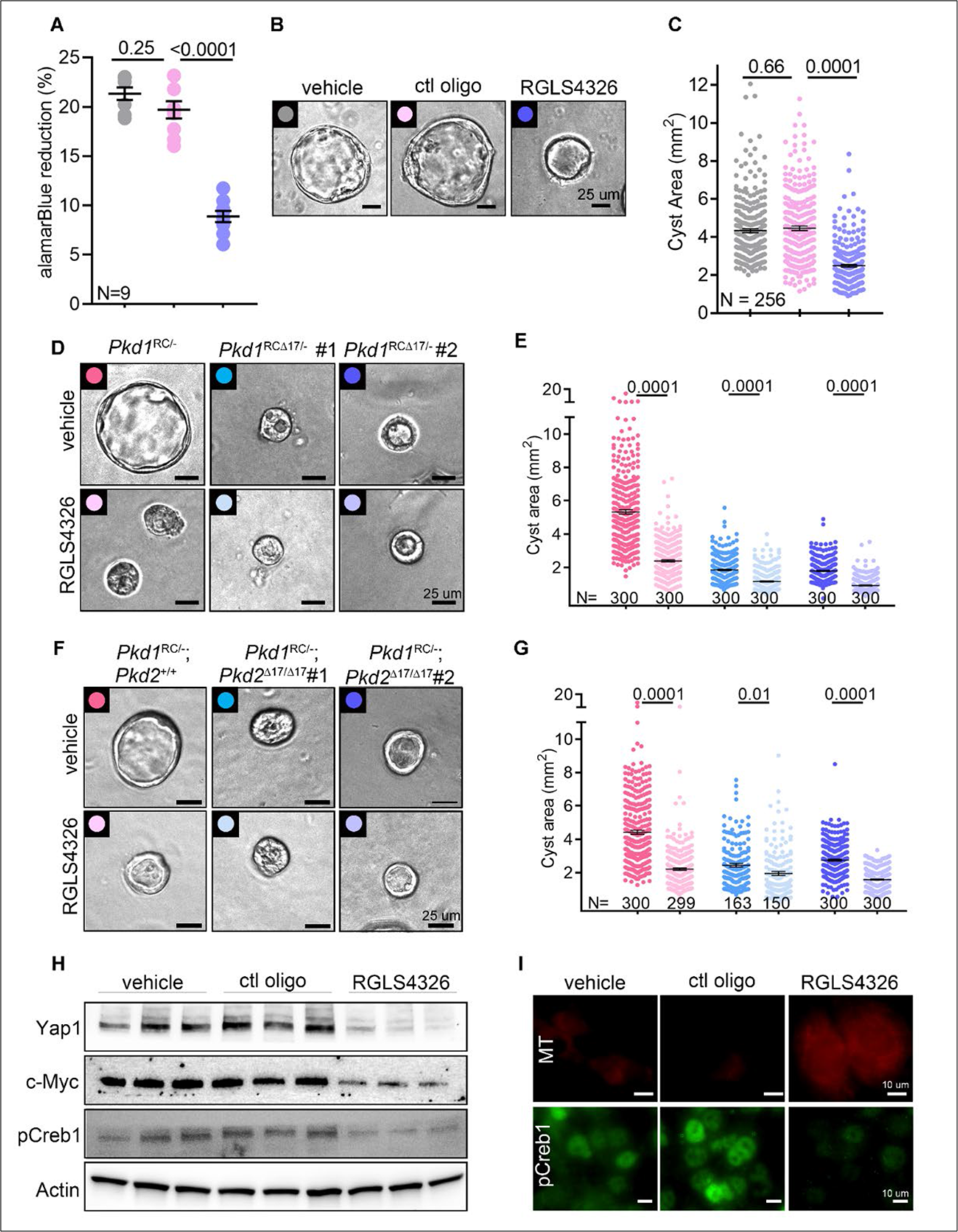
*Pkd1*^RC/-^ cells were transfected with 100 μM RGLS4326, 100 μM control oligonucleotide, or vehicle control. Seventy-two hours later, cells were equivalently seeded in 96-well plates for 12 hours for the alamarBlue assay or placed in Matrigel for seven days for a 3D cyst assay. **A.** Reduced proliferation of *Pkd1*^RC/-^ cells treated with RGLS4326 compared to cells treated with control oligonucleotide or vehicle control. **B-C.** Representative images and quantification showing marked reduction in cyst size of RGLS4326-treated compared to vehicle or control oligonucleotide-treated *Pkd1*^RC/-^ cells. No significant change was observed in cyst size between the vehicle control and control oligonucleotide-treated groups. **D-E.** Representative images and quantification showing cyst size of vehicle- or RGLS4326-treated *Pkd1*^RC/-^*, Pkd1*^RCΔ17/-^ (clone#1) or *Pkd1*^RCΔ17/-^(clone#2). **F-G.** Representative images and quantification showing cyst size of vehicle- or RGLS4326-treated *Pkd1*^RC/-^, *Pkd1*^RC/-^; *Pkd2*^Δ17/Δ17^ (clone#1) or *Pkd1*^RC/-^; *Pkd2*^Δ17/Δ17^ (clone#2). **H.** Immunoblots showing PC1, PC2, Yap1, c-Myc, and pCreb1 expression in *Pkd1*^RC/-cells^ transfected with a vehicle, control oligonucleotide, or RGLS4326. **I.** MitoTracker labeling and anti-pCreb1 immunostaining in *Pkd1*^RC/-^ cells transfected with a vehicle, control oligonucleotide, or RGLS4326. Error bars indicate SEM. Statistical analysis: one-way ANOVA and Tukey’s multiple comparisons test (A, C, E, G).

**Supplementary Figure 11:**
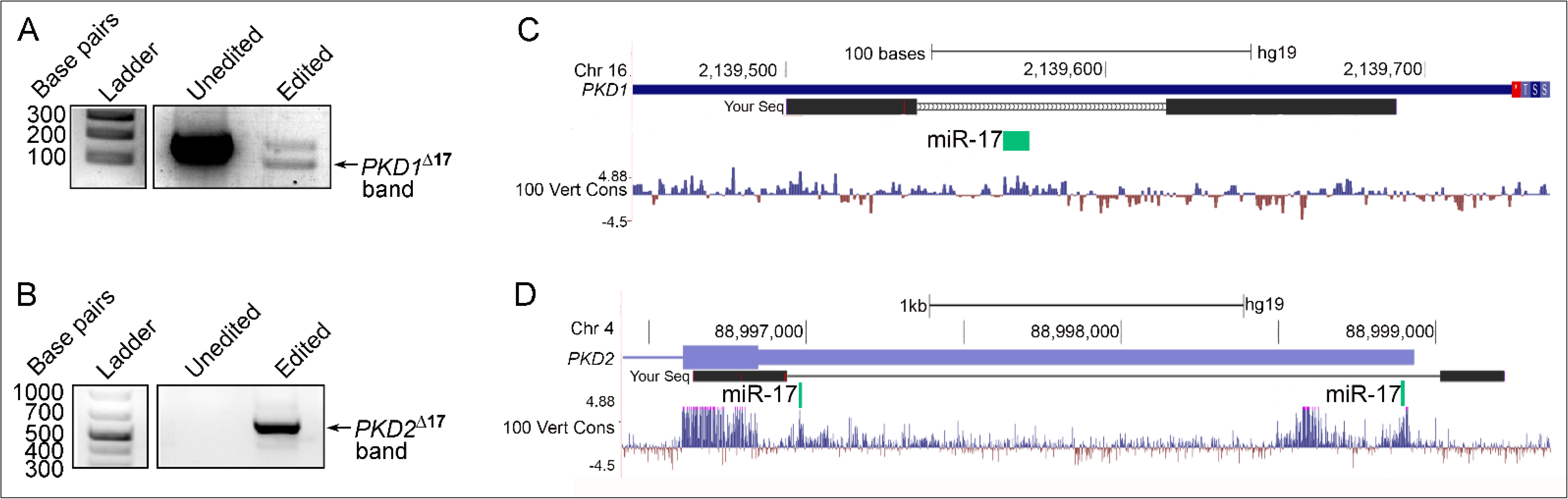
Genotyping of CRISPR-edited primary human ADPKD cultures. **A-B.** PCR products obtained after amplifying the DNA encoding the *PKD1* (**A**) or *PKD2* (**B**) 3’UTR segment from unedited parental and CRISPR-edited human ADPKD cultures. The arrows indicate the PCR bands resulting from miR17 motif deletion in the *PKD1* (**A**) and *PKD2* (**B**) genes. **C-D.** Sanger sequencing of the PCR product (black rectangles) aligned with the human genome (purple rectangles). The deleted region contains the miR-17 binding site (green rectangles).

**Supplementary Figure 12:**
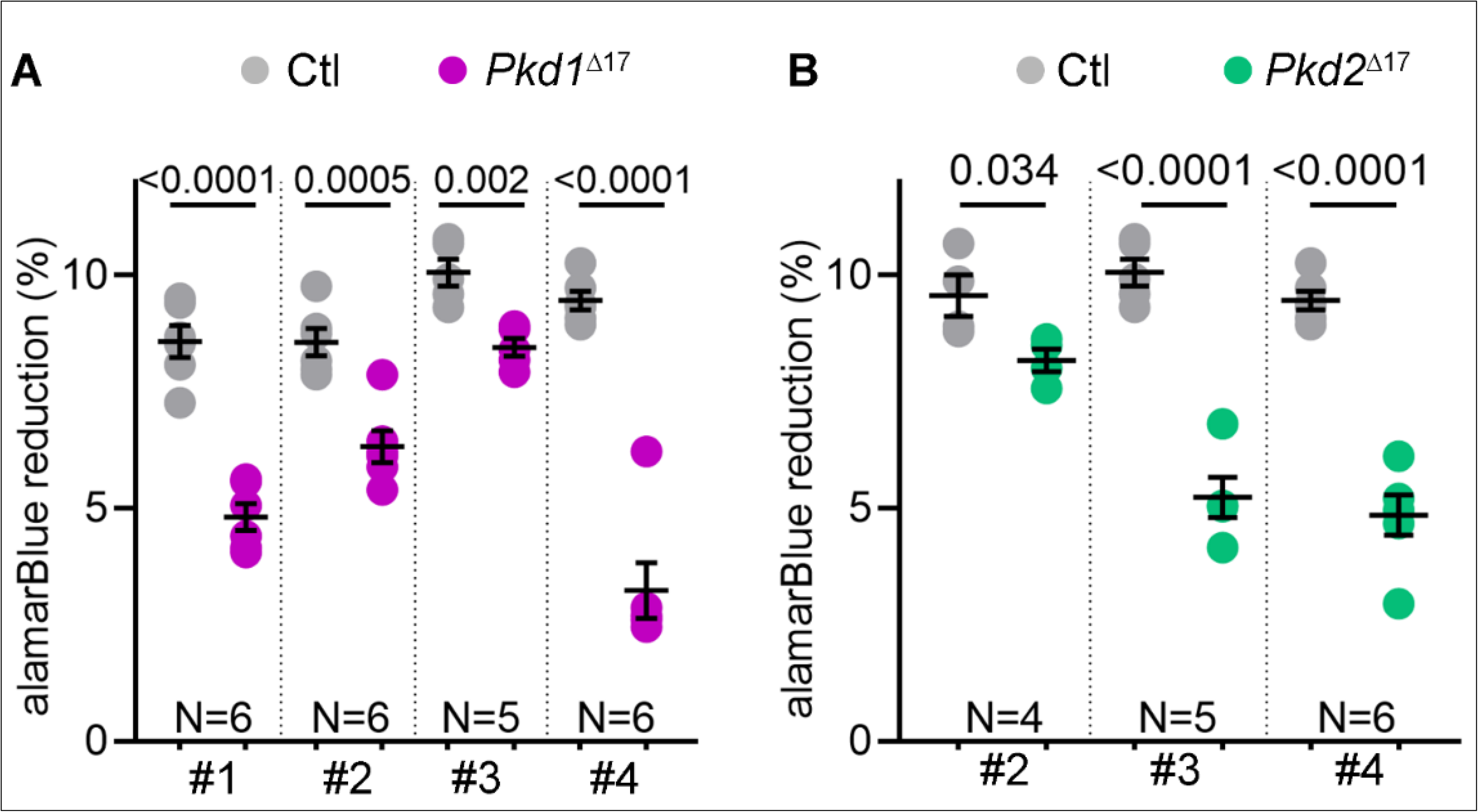
Reduced proliferation in *PKD1*^*Δ17-*^ and *PKD2*^*Δ17*^-edited ADPKD cultures. **A-B.** AlamarBlue-assessed proliferation of ADPKD donor cultures that were CRISPR edited to remove the miR-17 motif in either the *PKD1* (pink) or *PKD2* (green) gene compared to their respective unedited parental controls (Ctl, gray). Error bars indicate SEM. Statistical test: Student’s *t*-test.

