## Supplementary Figures for "*PKD1* and *PKD2* mRNA cis-inhibition drives polycystic kidney disease progression"

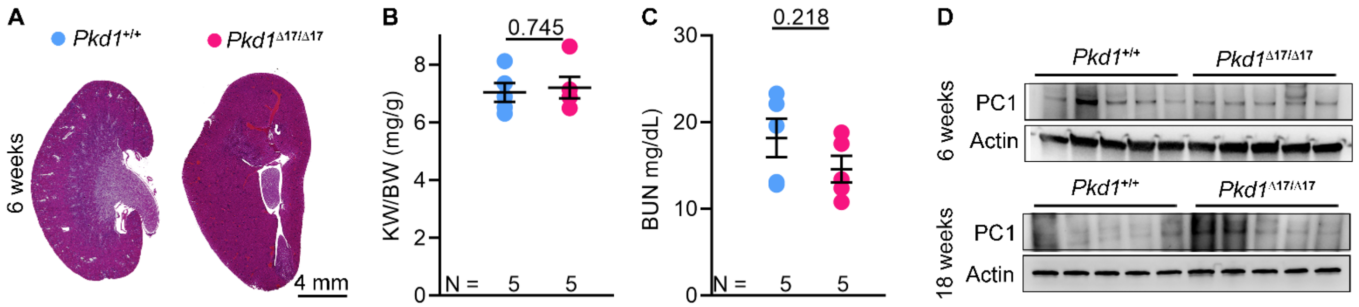


**Supplementary Figure 1: Characterization of *Pkd1*$^{\Delta17/\Delta17}$ mice. A.** H&E-stained kidney sections of 6-week-old *Pkd1*$^{+/+}$ and *Pkd1*$^{\Delta17/\Delta17}$ mice are shown. Both mice exhibited normal kidney histology. **B & C.** Kidney-weight-to-body-weight (KW/BW) ratio and BUN levels of 6-week-old *Pkd1*$^{+/+}$ and *Pkd1*$^{\Delta17/\Delta17}$ mice are shown. **D.** Immunoblot showing equivalent PC1 expression in kidneys of 6 or 18-week-old *Pkd1*$^{+/+}$ and *Pkd1*$^{\Delta17/\Delta17}$ mice. Error bars indicate SEM. Statistical analysis student's t-test (B-C).

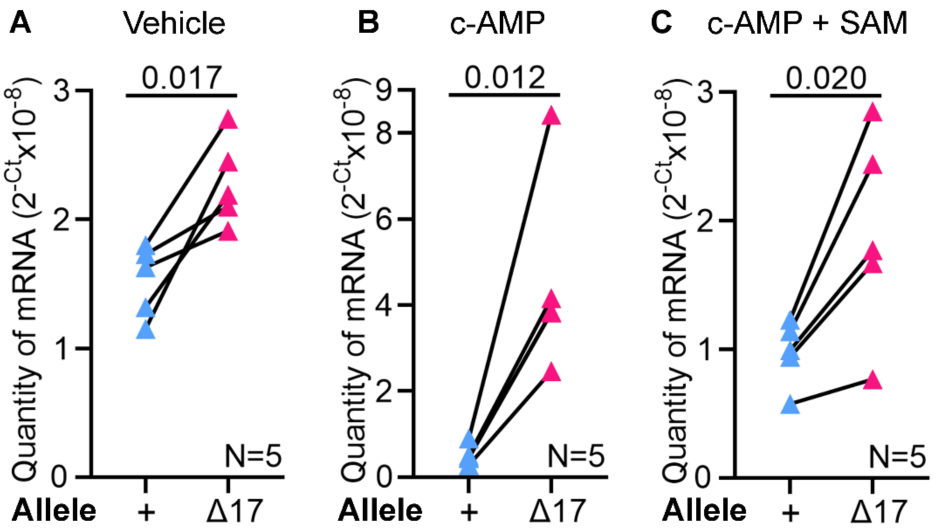


**Supplementary Figure 2: *Pkd1* is cis-inhibited via its miR-17 3'-UTR motif.** Allele-specific qRT-PCR analysis showing the quantity of *Pkd1* mRNAs produced by the wildtype (+) and Δ17 alleles in *ex vivo* kidney cultures of *Pkd1*$^{\Delta17/+}$ mice treated with vehicle (**A**), c-AMP (**B**), or c-AMP plus SAM (**C**). The *Pkd1*$^{\Delta17}$ allele produced more mRNA transcripts compared to the *Pkd1*$^{+}$ allele. This difference was even more pronounced in the presence of c-AMP. Statistical analysis: paired t-test. Error bars indicate SEM.

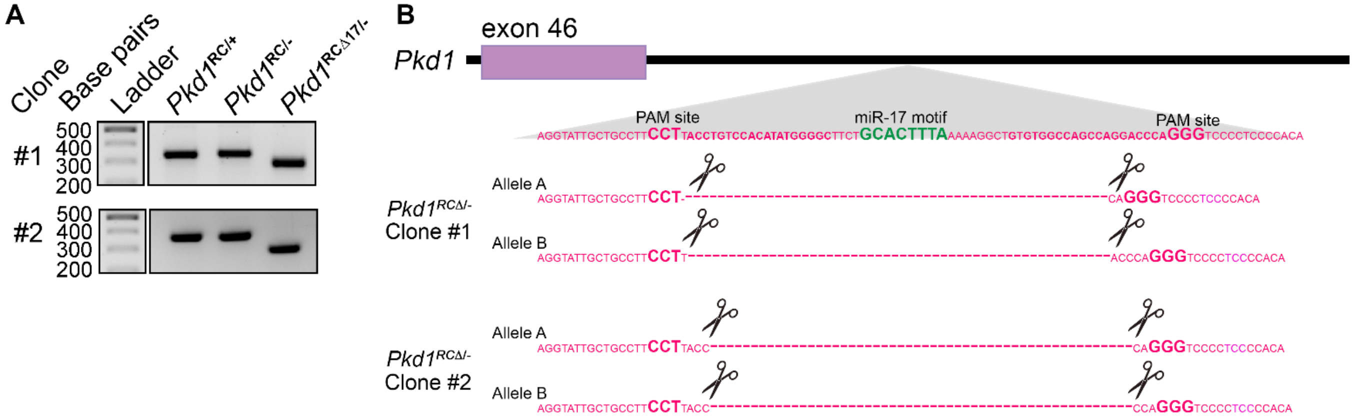


**Supplementary Figure 3: Characterization of CRISPR-edited *Pkd1*$^{RC/-}$ cell lines. A.** PCR products obtained after amplifying the DNA (encoding the *Pkd1* 3'UTR segment) from parental and CRISPR-edited cell lines. The lower band indicates the $\Delta 17$ genotype. **B.** Graphical illustration of Sanger sequencing results from the $\Delta 17$ bands of each clone confirm deletion of the miR-17 motif from both *Pkd1* alleles.

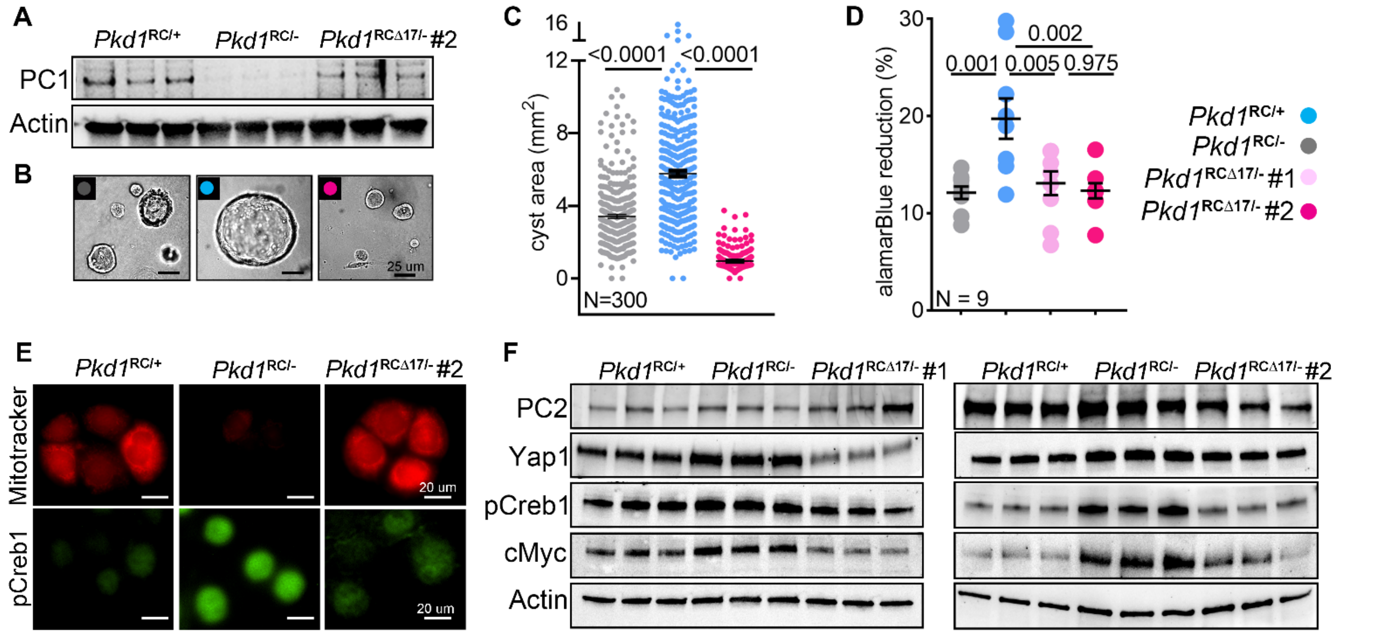


**Supplementary Figure 4. Phenotypic characterization of CRISPR-edited *Pkd1$^{RC\Delta17}$* cell lines. A.** Western blot showing PC1 derepression in *Pkd1*$^{RC\Delta17/-}$ clone #2 compared to *Pkd1*$^{RC/-}$ parent cell line. Actin is used as a loading control. **B, C.** 3D cyst images and quantification showing a substantial reduction in cyst size of *Pkd1*$^{RC\Delta17/-}$ cells compared to *Pkd1*$^{RC/-}$ cells. **D.** Alamarblue assay showing reduced proliferation of *Pkd1* clones which lack miR-17 motif at 12 hours compared to parent cell line. **E.** Mitotracker images and IF staining for pCreb1 showing restored mitochondrial membrane potential (red) and reduced pCreb1 (green) expression in *Pkd1*$^{RC\Delta17/-}$ clone #2 compared to parental cell line. **F.** Western blot characterization of both *Pkd1*$^{RC\Delta17/-}$ cell lines showing reduced expression of cyst promoting genes Yap1, pCreb1, c-Myc compared to parental *Pkd1*$^{RC/-}$ cells. As a pertinent control, PC2 expression remained unchanged. Actin serves as the loading control. Error bars indicate SEM. Statistical analysis: one-way ANOVA, Tukey's multiple comparisons test (C, D).

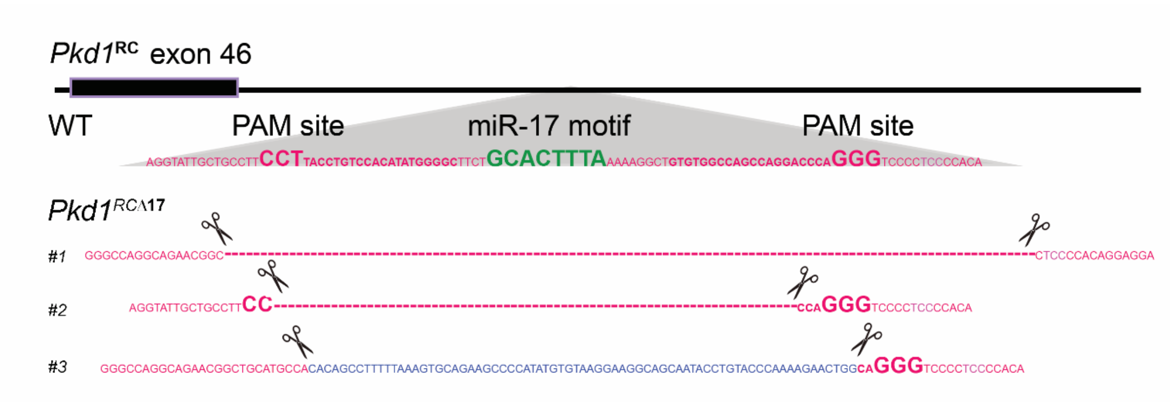


**Supplementary Figure 5**. **Characterization of CRISPR-edited *Pkd1*$^{RC/RC}$ mice**. Graphical illustration of Sanger sequencing from tail DNA from the three CRISPR-edited founders. Founders #1 and #2 harbor 108 bp and 53 base pair deletions, respectively, including the miR-17 motif. Founder #3 also lacked the miR-17 motif but acquired a 72 base pair insertion (blue), resulting in a net loss of 18 base pairs in the 3'-UTR sequence.

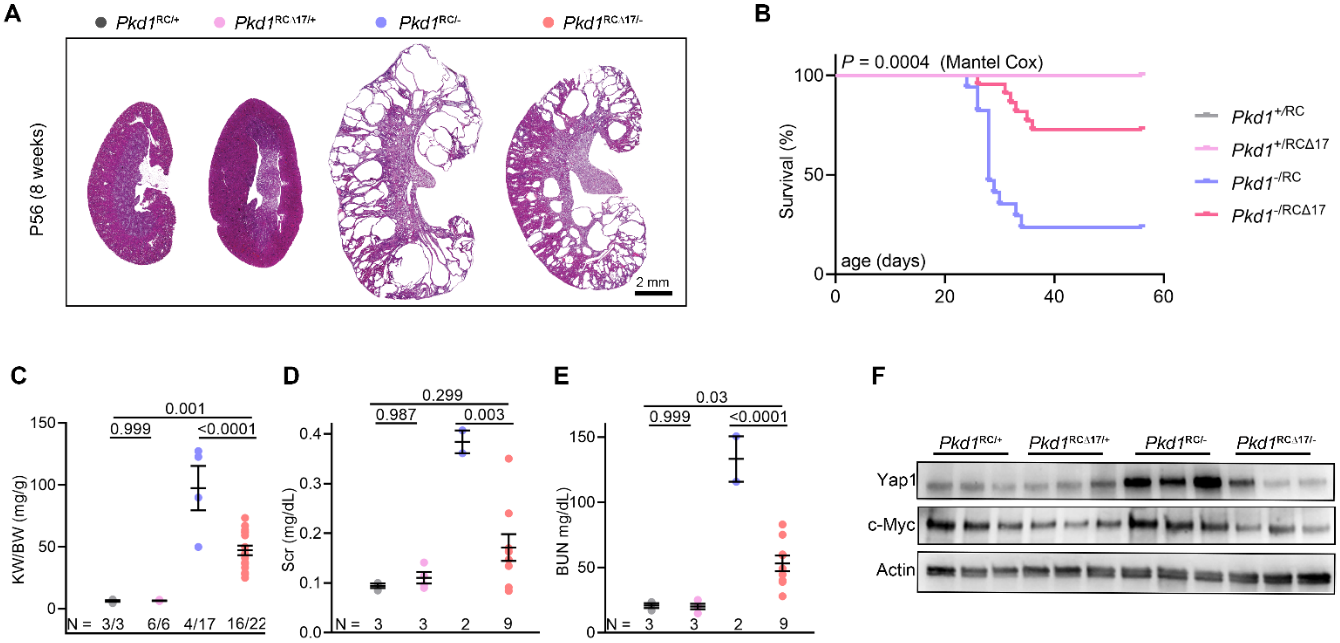


**Supplementary Figure 6: Monoallelic *Pkd1* derepression suppresses disease progression.** A cohort of progeny derived from founder #2 was prospectively monitored till 8-weeks of age. **A.** H&E-stained kidney sections of mice of the indicated genotypes that survived till 8-weeks of age are shown. **B.** Kaplan-Meir survival curves of mice with the indicated genotypes are shown. **C-E**. KW/BW, serum creatinine (Scr), and BUN levels of the surviving 8-week-old mice with indicated genotypes are shown. **F**. Immunoblots showing Yap1 and c-Myc expression in kidneys of 18-day-old mice with the indicated genotypes. Actin serves as the loading control. Error bars indicate SEM. Statistical analysis: one-way ANOVA, Tukey's multiple comparisons test (C-E); Log-rank Mantel-Cox (B).

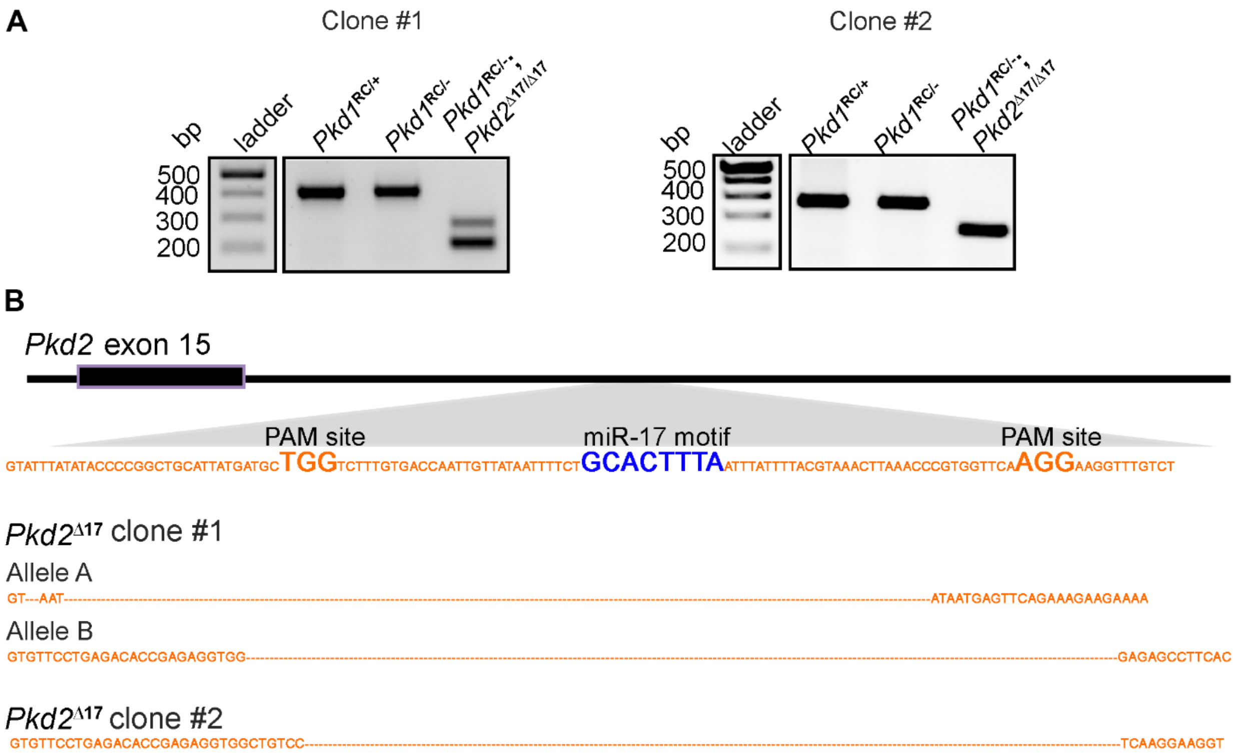


**Supplementary Figure 7: Characterization of *Pkd1*$^{RC/-}$ cell lines lacking miR-17 motif from *Pkd2* 3'-UTR. A.** PCR products obtained after amplifying the DNA (encoding the *Pkd2* 3'UTR segment) from parental and CRISPR-edited cell lines. The lower bands indicate the Δ17 genotype. **B.** Graphical illustration of Sanger sequencing results from the Δ17 bands of each clone confirm deletion of the miR-17 motif from both *Pkd2* alleles.

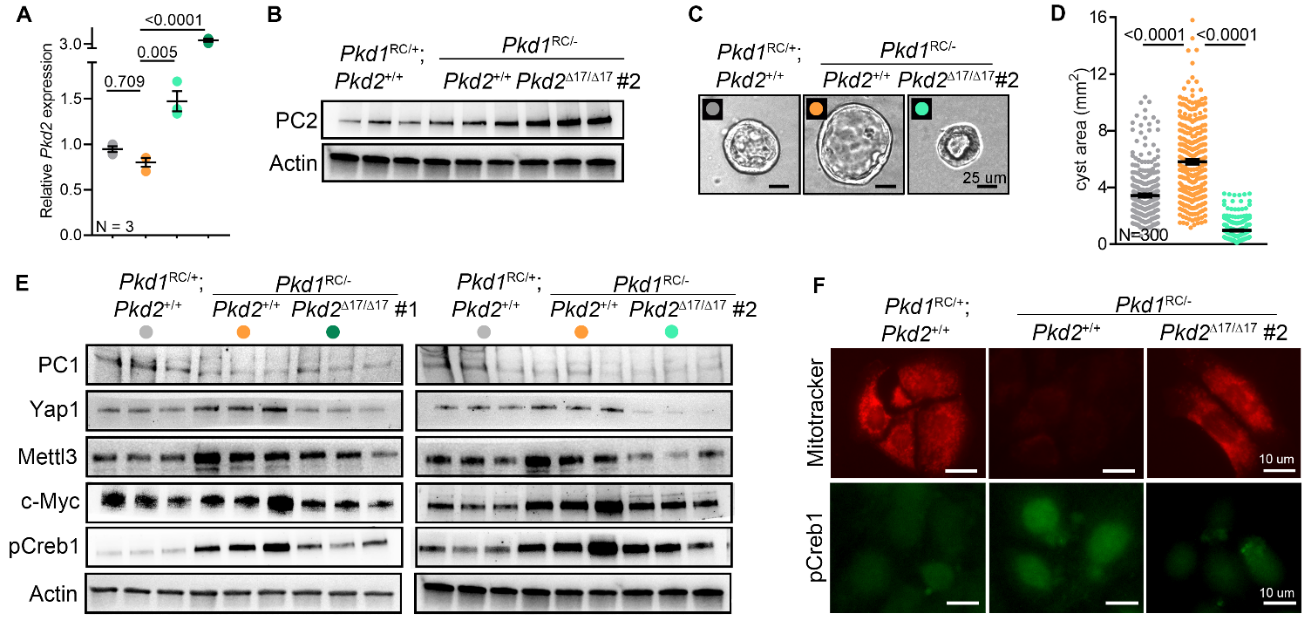


**Supplemental Figure 8. Phenotypic characterization of CRISPR-edited *Pkd1*$^{RC/-}$*; Pkd2*$^{\Delta 17/\Delta 17}$ cell lines.** **A, B.** qRT-PCR and Western blot showing *Pkd2* and Polycystin-2 derepression, respectively, in both *Pkd1*$^{RC/-}$*; Pkd2*$^{\Delta 17/\Delta 17}$ clones #1 and #2 compared to the parental *Pkd1*$^{RC/-}$ cell line. **C-D** Representative images and quantification of 3D cyst assay comparing indicated cell lines showing marked reduction in cyst size of *Pkd1*$^{RC/-}$*; Pkd2*$^{\Delta 17/\Delta 17}$ clone #2 compared to parent *Pkd1*$^{RC/-}$ cell line. **E.** Western blot characterization showing reduced expression of cyst promoting genes Yap1, Mettl3, c-Myc, and pCreb1 in both *Pkd1*$^{RC/-}$*; Pkd2*$^{\Delta 17/\Delta 17}$ clones compared to the parent *Pkd1*$^{RC/-}$ cell line. Pertinently, PC1 expression remained unchanged. Actin serves as a loading control. **F.** Mitotracker images and IF staining for pCreb1 showing restored mitochondrial membrane potential (red) and reduced pCreb1 (green) expression in *Pkd1*$^{RC/-}$*; Pkd2*$^{\Delta 17/\Delta 17}$ clone #2 compared to parental cell line. Error bars indicate SEM. Statistical analysis: one-way ANOVA, Tukeys' multiple comparisons test (A, D).

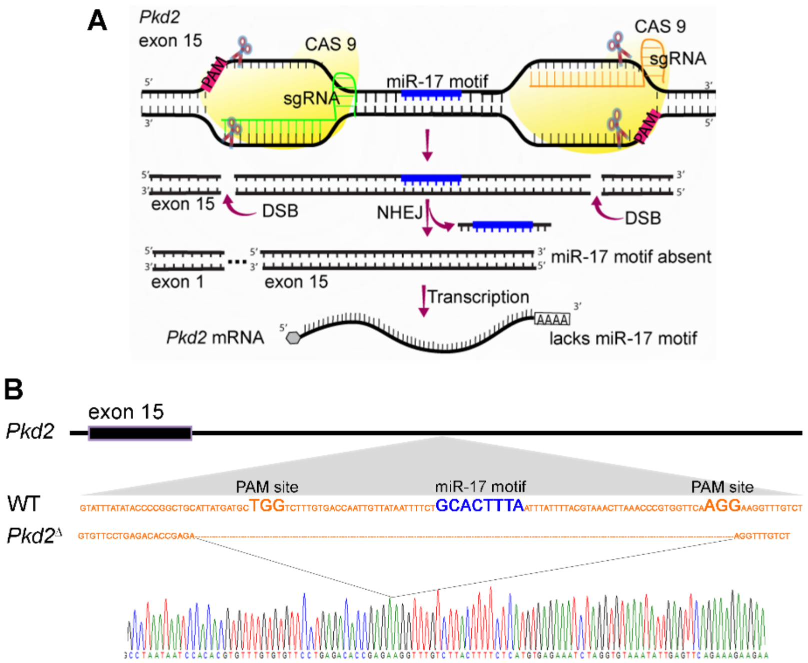


**Supplementary Figure 9**. **CRISPR-editing and characterization of** *Pkd2*$^{\Delta 17}$ **mice that lack miR-17 motif in *Pkd2* 3'-UTR. A.** Graphic illustrates the CRISPR-editing strategy used to remove the miR-17 motif from exon 15 of *Pkd2*. **B**. Graphical illustration and Sanger sequencing results of PCR product from the tail DNA of founder mouse showing 139 base pair deletion from the *Pkd2* 3'-UTR, including the miR-17 motif.

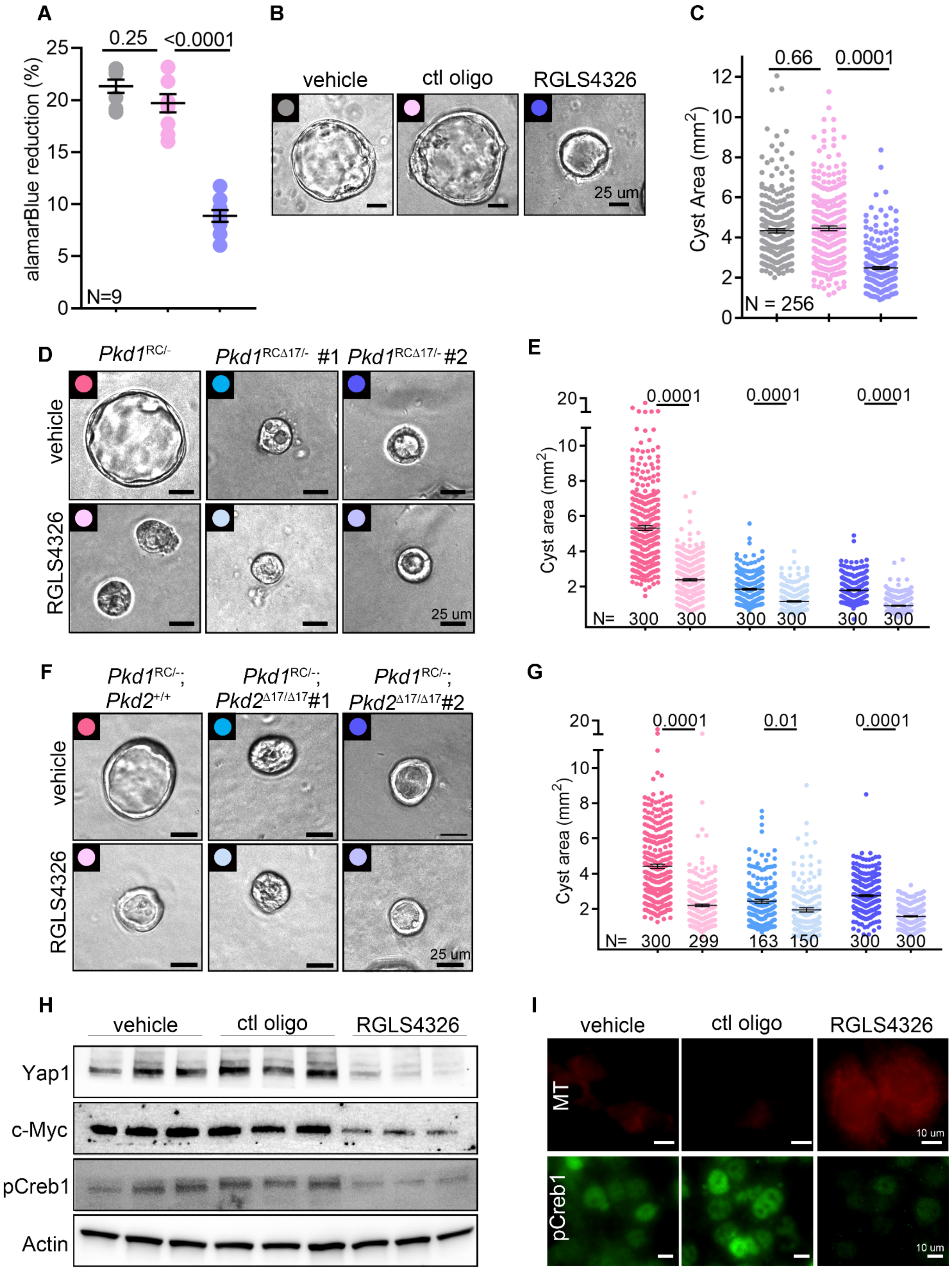
A
alamarBlue reduction (%)
0.25
<0.0001
N=9
B
vehicle
ctl oligo
RGLS4326
25 um
C
Cyst Area (mm2)
0.66
0.0001
N = 256
D
Pkd1RC/-
Pkd1RCΔ17/- #1
Pkd1RCΔ17/- #2
vehicle
RGLS4326
25 um
E
Cyst area (mm2)
0.0001
0.0001
0.0001
N= 300 300 300 300 300 300
F
Pkd1RC/-; Pkd2+/+
Pkd1RC/-; Pkd2Δ17/Δ17#1
Pkd1RC/-; Pkd2Δ17/Δ17#2
vehicle
RGLS4326
25 um
G
Cyst area (mm2)
0.0001
0.01
0.0001
N= 300 299 163 150 300 300
H
vehicle
ctl oligo
RGLS4326
Yap1
c-Myc
pCreb1
Actin
I
vehicle
ctl oligo
RGLS4326
MT
pCreb1
10 um

**Supplementary Figure 10:** *Pkd1*$^{RC/-}$ cells were transfected with 100 uM RGLS4326, 100 uM control oligonucleotide, or vehicle control. Seventy-two hours later, cells were equivalently seeded in 96 well plate for 12 hours for the alamarBlue assay or placed in matrigel for seven days for a 3D cyst assay. **A.** Reduced proliferation of *Pkd1*$^{RC/-}$ cells treated with RGLS4326 compared to cells treated with control oligonucleotide or vehicle control. **B-C.** Representative images and quantification showing marked reduction in cyst size of RGLS4326-treated compared to vehicle or control oligonucleotide-treated *Pkd1*$^{RC/-}$ cells. No significant change was observed in cyst size between vehicle control and control oligonucleotide-treated groups. **D-E.** Representative images and quantification showing cyst size of vehicle or RGLS4326-treated *Pkd1*$^{RC/-}$, *Pkd1*$^{RC\Delta17/-}$ (clone#1) or *Pkd1*$^{RC\Delta17/-}$(clone#2). **F-G**. Representative images and quantification showing cyst size of vehicle or RGLS4326-treated *Pkd1*$^{RC/-}$, *Pkd1*$^{RC/-}$; *Pkd2*$^{\Delta17/\Delta17}$ (clone#1) or *Pkd1*$^{RC/-}$; *Pkd2*$^{\Delta17/\Delta17}$ (clone#2). **H.** Immunoblots showing PC1, PC2, Yap1, c-Myc, and pCreb1 expression in *Pkd1*$^{RC/-}$ transfected with a vehicle, control oligonucleotide, RGLS4326. **I.** Mitotracker labeling and anti-pCreb1 immunostaining in *Pkd1*$^{RC/-}$ transfected with a vehicle, control oligonucleotide, or RGLS4326. Error bars indicate SEM. Statistical analysis: one-way ANOVA, Tukeys' multiple comparisons test (A, C, E, G).

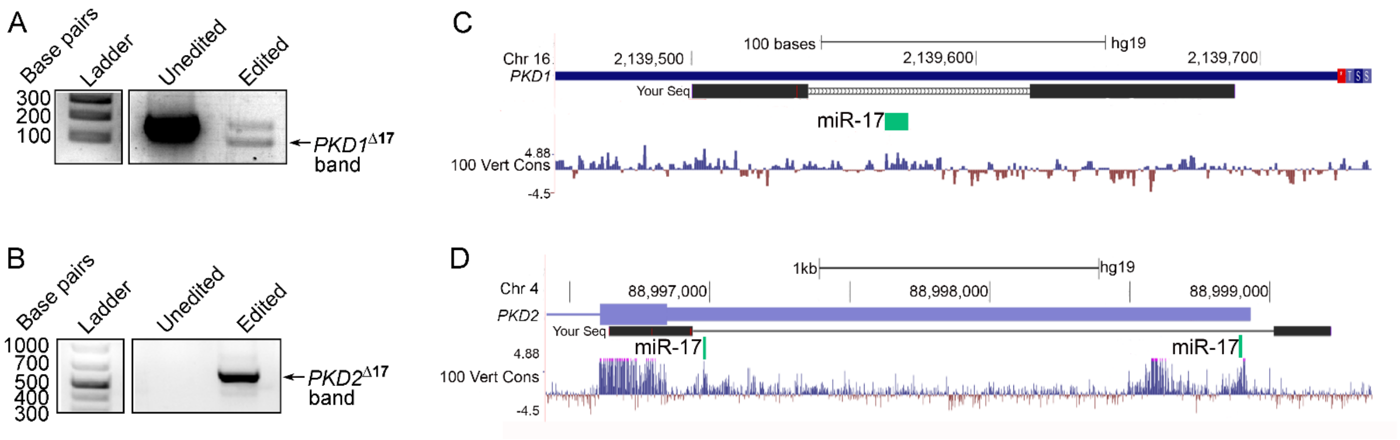


**Supplementary Figure 11: Genotyping of CRISPR-edited primary human ADPKD cultures. A-B.** PCR products obtained after amplifying the DNA encoding the *PKD1* (**A**) or *PKD2* (**B**) 3'UTR segment from unedited parental and CRISPR-edited human ADPKD cultures. The arrows indicate the PCR bands resulting from miR17 motif deletion in *PKD1* (**A**) and *PKD2* (**B**) genes. **C-D.** Sanger sequencing of the PCR product (black rectangles) aligned with the human genome (purple rectangles). The deleted region contains the miR-17 binding site (green rectangles).

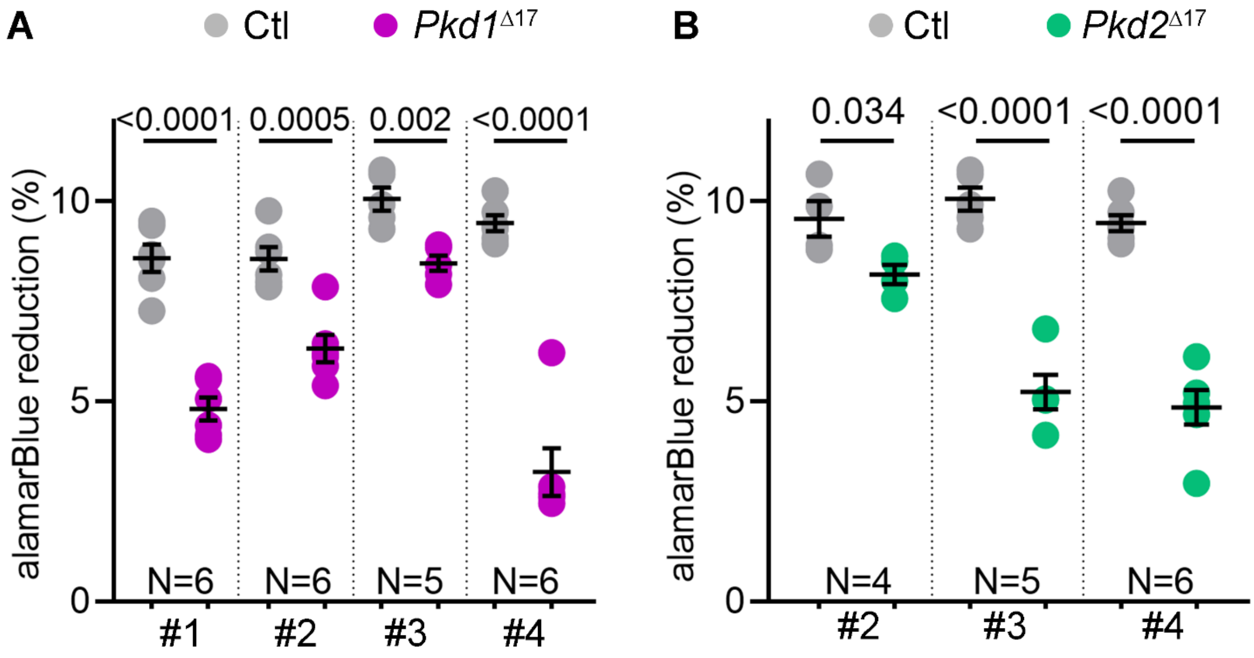


**Supplementary Figure 12: Reduced proliferation in *PKD1*$^{\Delta17}$ and *PKD2*$^{\Delta17}$ edited ADPKD cultures**. **A-B.** AlamarBlue-assessed proliferation of ADPKD donor cultures that were CRISPR-edited to remove the miR-17 motif in either the *PKD1* (pink) or *PKD2* (Green) gene compared to their respective unedited parental controls (Ctl, grey). Error bars indicate SEM. Statistical test: students *t*-test.
