## Supplementary Tables for "*PKD1* and *PKD2* mRNA cis-inhibition drives polycystic kidney disease progression"

| Cell line | $Pkd1^{RC/-}$ | | $Pkd1^{RC\Delta17\#1/-}$ | | $Pkd1^{RC/-}$ | | $Pkd1^{RC\Delta17\#1/-}$ | | $Pkd1^{RC/-}$ | | $Pkd1^{RC\Delta17\#1/-}$ | |
|---|---|---|---|---|---|---|---|---|---|---|---|---|
| **Stressor** | Ctl | cAMP | Ctl | cAMP | Ctl | Gluc | Ctl | Gluc | Ctl | SAM | Ctl | SAM |
| **Replicate** | | | | | | | | | | | | |
| 1 | 40.19 | 46.67 | 30.04 | 28.88 | 38.07 | 49.99 | 27.39 | 27.35 | 38.58 | 47.47 | 26.87 | 29.91 |
| 2 | 39.75 | 45.56 | 25.8 | 25.63 | 37.34 | 49.78 | 31.51 | 27.74 | 39.09 | 46.37 | 25.20 | 26.70 |
| 3 | 40.32 | 41.16 | 26.19 | 26.97 | 38.19 | 47.97 | 28.27 | 27.03 | 39.29 | 42.01 | 27.25 | 28.02 |
| 4 | 37.42 | 42.82 | 27.03 | 27.97 | 37.78 | 44.98 | 27.62 | 26.04 | 37.77 | 43.66 | 28.09 | 29.01 |
| 5 | 38.77 | 40.43 | 27.26 | 28.99 | 34.85 | 43.70 | 27.25 | 28.45 | 37.33 | 41.29 | 28.32 | 30.03 |
| 6 | 36.75 | 42.37 | 27.79 | 28.19 | 38.74 | 43.92 | 25.34 | 28.74 | 40.58 | 43.21 | 28.85 | 29.24 |
| 7 | 37.98 | 44.11 | 25.84 | 26.91 | 40.26 | 46.17 | 25.54 | 29.29 | 39.85 | 44.93 | 26.91 | 27.97 |
| 8 | 39.15 | 41.98 | 27.92 | 22.64 | 38.77 | 48.13 | 28.19 | 29.96 | 39.51 | 42.83 | 28.98 | 23.72 |
| ***P* value** | | | | | | | | | | | | |
| Ctl vs Stressor | 0.0002 | | | | <0.0001 | | | | <0.0001 | | | |
| Ctl vs Stressor | | | .9952 | | | | .7041 | | | | .9296 | |
| Ctl vs Ctl | <0.0001 | | | | <0.0001 | | | | <0.0001 | | | |

**Supplementary Table 1. AlamarBlue Assay on *Pkd1*$^{RC/-}$ and *Pkd1*$^{RC\Delta17\#1/-}$ cells.** Raw values of percentage reduction are shown for each biological replicate. The results of one-way ANOVA followed by post-hoc Tukey's multiple comparisons test are listed.

| Cell line | ***Pkd1***$^{RC/-}$ | | ***Pkd1***$^{RC/-}$ ***Pkd2***$^{\Delta17/\ \Delta17\#1}$ | | ***Pkd1***$^{RC/-}$ | | ***Pkd1***$^{RC/-}$ ***Pkd2***$^{\Delta17/\ \Delta17\#1}$ | | ***Pkd1***$^{RC/-}$ | | ***Pkd1***$^{RC/-}$ ***Pkd2***$^{\Delta17/\ \Delta17\#1}$ | |
|---|---|---|---|---|---|---|---|---|---|---|---|---|
| **Stressor** | Ctl | cAMP | Ctl | cAMP | Ctl | Gluc | Ctl | Gluc | Ctl | SAM | Ctl | SAM |
| **Replicate** | | | | | | | | | | | | |
| 1 | 32.32 | 36.36 | 18.7 | 17.73 | 27.94 | 34.09 | 23.44 | 21.66 | 38.11 | 50.04 | 29.66 | 31.93 |
| 2 | 32.14 | 36.15 | 17.51 | 18.6 | 28.8 | 34.05 | 21.61 | 21.57 | 43.64 | 51.71 | 29.57 | 30.41 |
| 3 | 30.55 | 36.5 | 17.79 | 19 | 29.57 | 34.49 | 22.48 | 19.47 | 42.4 | 50.53 | 31.2 | 30.42 |
| 4 | 30.43 | 37.4 | 19.85 | 17.84 | 29.49 | 34.34 | 20.42 | 20.87 | 43.66 | 51.62 | 30.76 | 32.56 |
| 5 | 31.35 | 37.14 | 19.25 | 19.67 | 28.05 | 35.09 | 18.43 | 21.22 | 43.51 | 48.1 | 30.62 | 32 |
| 6 | 32.44 | 38.96 | 19.17 | 19.8 | 27.4 | 35.16 | 19.93 | 21.22 | 43.83 | 53.31 | 30.21 | 32.06 |
| 7 | 32.28 | 36.35 | 19.06 | 21.36 | 28.29 | 35.84 | 20.94 | 18.87 | 42.98 | 49.38 | 30.31 | 30.7 |
| 8 | 32.55 | 38.71 | 18.34 | 17.82 | 27.86 | 36.1 | 19.45 | 20.13 | 43.6 | 48.67 | 33.71 | 30.95 |
| ***P* value** | | | | | | | | | | | | |
| Ctl vs Stressor | <0.0001 | | | | <0.0001 | | | | <0.0001 | | | |
| Ctl vs Stressor | | | .9519 | | | | .9811 | | | | .8421 | |
| Ctl vs Ctl | <0.0001 | | | | <0.0001 | | | | <0.0001 | | | |

**Supplementary Table 2. AlamarBlue Assay on *Pkd1*$^{RC/-}$ and *Pkd1*$^{RC/-}$;*Pkd2*$^{\Delta17/\ \Delta17\#1}$ cells.** Raw values of percentage reduction are shown for each biological replicate. The results of one-way Anova followed by post-hoc Tukey's multiple comparisons test are listed.

| | SgRNA sequence (5'-3') | | Genotyping Primers (5'-3') | |
|---|---|---|---|---|
| | **Upstream** | **Downstream** | **Forward Primer** | **Reverse Primer** |
| ***Pkd1*** | GCCCATATGTGGACAGGTA | GTGTGGCCAGCCAGGACCCA | CTAGGGGTCTTGGCCATTCC | CTGAACCTGAGGACTTGGGG |
| ***Pkd2*** | CGAACTTATCATTGTCGTAC | AAACTTAAACCCGTGGTTCA | GTTCCTGAGACACCGAGAGG | GGCAGCAAATCAGCGTACTT |
| ***PKD1*** | ACGTAGGTTCCCCAGAGAGCA | TGTGTGGCCAACCAGGACCC | TGAGGACTCGGGGAAATAAA | CGTGGAGTCGGAGTGGAC |
| ***PKD2*** | TTATATGCCCTGACCACCAT | AATGTTTTCGGACATAGAGA | GGTCGTGACAGTGAAATCCA | (WT) CAGAAGATTAGCAATCGGTCAC (Deletion Specific) TGGCAGCCTATGCTGTTATG |

**Supplementary Table 3. SgRNA Sequences and Genotyping Primer Sequences**

| | Forward (5'-3') | Reverse (5'-3') |
|---|---|---|
| *Pkd1* | CTAGACCTGTCCCACAACCTA | GCAAACACGCCTTCTTCTAATGT |
| *Pkd2* | GCGTGGTACCCTCTTGGCAGTT | CACGACAATCACAACATCC |
| *Pkd1* (+ allele) | CATATGGGGCTTCTGCACTT | GAGGCTGGGTACTCACTTGG |
| *Pkd1* (Δ allele) | ATTGCTGCCTTCCTTACCCC | CTGGGTACTCACTTGGTCCA |

**Supplementary Table 4. Q-PCR primer sequences**
